# Construction of a third NAD^+^ *de novo* biosynthesis pathway

**DOI:** 10.1101/2020.10.18.342451

**Authors:** Yong Ding, Xinli Li, Geoff P. Horsman, Pengwei Li, Min Wang, Jine Li, Zhilong Zhang, Weifeng Liu, Bian Wu, Yong Tao, Yihua Chen

## Abstract

Only two *de novo* biosynthetic routes to nicotinamide adenine dinucleotide (NAD^+^) have been described, both of which start from a proteinogenic amino acid and are tightly controlled. Here we establish a C3N pathway starting from chorismate in *Escherichia coli* as a third NAD^+^ *de novo* biosynthesis pathway. Significantly, the C3N pathway yielded extremely high cellular concentrations of NAD(H) in *E. coli*. Its utility in cofactor engineering was demonstrated by introducing the four-gene C3N module to cell factories to achieve higher production of 2,5-dimethylpyrazine and develop an efficient C3N-based whole-cell bioconversion system for preparing chiral amines. The wide distribution and abundance of chorismate in most kingdoms of life implies a general utility of the C3N pathway for modulating cellular levels of NAD(H) in versatile organisms.

## Introduction

Nicotinamide adenine dinucleotide (NAD^+^) is a universally essential central metabolite consisting of adenosine monophosphate (AMP) linked to nicotinamide mononucleotide (NMN). Since its discovery as the first ‘cozymase’ over 100 years ago, two *de novo* biosynthesis pathways for NAD^+^ have been reported (Fig. 1)^1^. Both pathways use a proteinogenic amino acid as the precursor for nicotinamide and converge at a common intermediate quinolinic acid (QA). In plants and most bacteria, the *de novo* biosynthesis of NAD^+^ starts from L-aspartate (L-Asp), which is converted to QA by L-Asp oxidase (NadB) and quinolinate synthase (NadA) (Pathway I); in mammals, fungi, and some bacteria, nicotinamide is derived from conversion of L-tryptophan (L-Trp) to 3-hydroxyanthranilic acid (3-HAA) by four reactions of the kynurenine pathway (Pathway II). Oxidation of 3-HAA by 3-HAA 3,4-dioxygenase yields 2-amino-3-carboxymuconate semialdehyde (ACMS), which undergoes spontaneous ring closure to form QA^1–3^. The QA generated in both pathways is converted to NAD^+^ *via* a common three-enzyme pathway (catalyzed by enzymes NadC-E): phosphoribosylation of QA to generate nicotinic acid mononucleotide, AMP addition to form nicotinic acid adenine dinucleotide, and a final amidation to afford NAD^+^. In addition to the two *de novo* biosynthesis pathways, NAD^+^ is also regenerated by different salvage pathways using varied pyridines (nicotinic acid, nicotinamide, and nicotinamide riboside) as precursors^1, 4^.

**Fig. 1.**
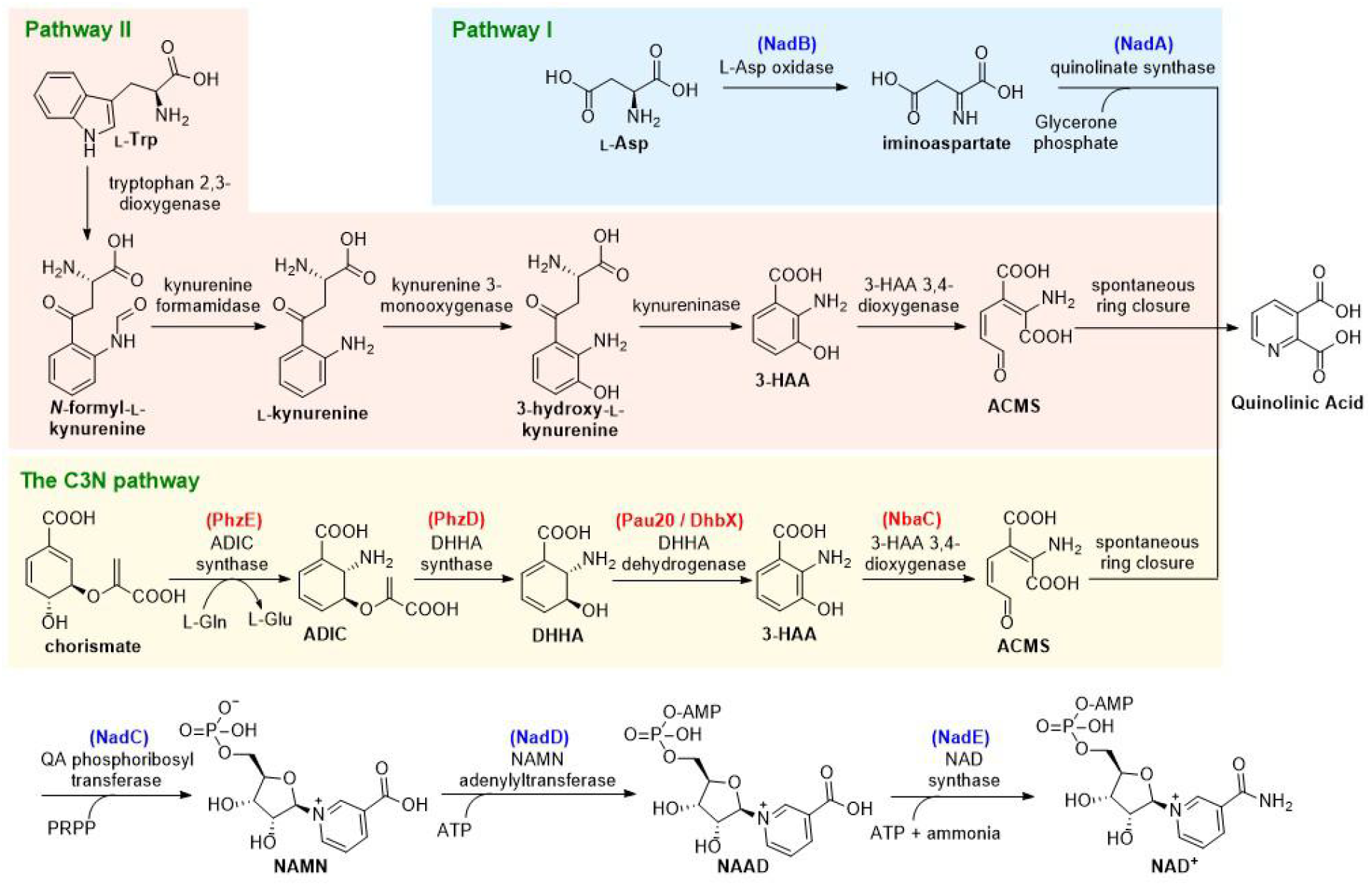
Three NAD^+^ *de novo* biosynthesis pathways. The enzymes used in this work are in parenthesis and color-coded as either red (from secondary metabolic processes like natural product biosynthesis and aromatic compound degradation) or blue (from NAD^+^ biosynthesis). ADIC: 2-amino-2-deoxyisochorismate; DHHA: 2,3-dihydro-3-hydroxyanthranilic acid; ACMS: 2-amino-3-carboxymuconate semialdehyde; 3-HAA: 3-hydroxyanthranilic acid; PRPP: 5-phosphoribosyl diphosphate; NAMN: nicotinic acid mononucleotide; NAAD: nicotinic acid adenine dinucleotide.

NAD^+^ and its reduced form NADH serve as hydride acceptor and donor, respectively, in thousands of redox reactions in vital metabolic pathways such as glycolysis, citric acid cycle, fatty acid degradation, and oxidative phosphorylation^5^. Although cellular levels of NAD(H) are usually sufficient to support NAD(H)-dependent oxidoreductases under physiological conditions, NAD(H) concentrations may become rate-limiting when these enzymes are overproduced in engineered cell factories^6^. Indeed, many such examples of limiting NAD(H) have been documented since the first report of *Escherichia coli* cells overexpressing *Vibrio harveyi* luciferase in 1982^7, 8^. NAD(H) deficiency is also medically significant; for example carcinoid syndrome patients, in whom L-Trp is depleted by excessive production of serotonin, exhibit pellagra-like symptoms caused by NAD(H) deficiency^9^.

NAD^+^ has also been recently identified as the substrate for NAD^+^-consuming enzymes involved in signal transduction pathways that regulate crucial biological processes including DNA repair, transcription, and cell cycle progression^10–12^. In some cases, cellular levels of NAD(H) are reduced dramatically by NAD^+^-consuming enzymes; for example, the hyper-activation of poly(ADP-ribose) polymerase-1 reduced the NAD^+^ pool by about 80%^11^. Consequently, increasing cellular NAD(H) concentrations can stimulate the NAD^+^-consuming enzymes and modulate downstream biological processes. For example, increasing NAD(H) levels can activate sirtuins and improve mitochondrial homeostasis and extend the lifespan of different species^13, 14^.

Several strategies have been developed to enhance the catalytic efficiencies of NAD(H)-dependent or NAD^+^-consuming enzymes by increasing cellular NAD(H) levels. These include supplementation with NAD^+^ or its pyridine precursors, limiting NAD^+^ consumption, and reinforcing NAD^+^ salvage or *de novo* biosynthesis pathways^15–17^. However, each strategy has its limitations: supplementation is restricted by poor cellular uptake of NAD^+^; reducing NAD^+^ consumption and accelerating its recycling *via* the salvage pathways can only replenish but cannot expand cellular NAD(H) pools. Considering that the highest cellular NAD(H) levels are limited in large part by NAD^+^ *de novo* biosynthetic capacity, manipulating these *de novo* pathways should efficiently expand cellular NAD(H) pools. The most frequently used strategy for increasing cellular NAD(H) levels is by reinforcing the NAD^+^ salvage pathways instead of manipulating *de novo* biosynthesis. However, the few reported attempts to manipulate NAD^+^ *de novo* pathways yielded only very limited^18^ or nonexistent increases in cellular NAD(H) levels^19^, probably due to the stringent regulation of NAD^+^ *de novo* biosynthesis at transcriptional^20^, translational^21^, and post-translational levels^22^. In addition, because both NAD^+^ *de novo* biosynthesis pathways start from a proteinogenic amino acid, their activation may deplete amino acid pools required to efficiently overproduce proteins in engineered cell factories.

In this study, we decoupled NAD^+^ *de novo* biosynthesis and protein synthesis by designing a C3N pathway as the third NAD^+^ *de novo* biosynthesis pathway, which uses chorismate as the precursor for nicotinamide. The C3N pathway was constructed in *E. coli* by combining genes from secondary metabolism with the latter steps of pathway I. It effectively circumvents the tight regulatory controls on NAD^+^ *de novo* biosynthesis and enables extremely high cellular concentrations of NAD(H) in the recombinant *E. coli* strains. We genetically packaged the C3N pathway as a four-gene C3N module for plug-and-play installation to significantly boost cellular NAD(H) concentrations in *E. coli*. Its utility in cofactor engineering was demonstrated by improving the bioconversion efficiency of 2,5-dimethylpyrazine (DMP) and by developing a C3N-based whole-cell system for efficient production of chiral amines.

## Results and Discussion

### Conceptual design of a third NAD^+^ *de novo* biosynthesis pathway

The conceptual basis for an alternative NAD^+^ *de novo* biosynthesis pathway arose from the observation that several secondary metabolites contain structures derived from 3-HAA^23–28^ (Supplementary Fig. 1a), which also occurs in primary metabolism as a key intermediate in NAD^+^ *de novo* biosynthesis pathway II (Fig. 1). The biosynthetic gene clusters encoding these natural products indicate that chorismate is converted to 3-HAA by three sequential reactions catalyzed by 2-amino-2-deoxyisochorismate (ADIC) synthase, 2,3-dihydro-3-hydroxyanthranilic acid (DHHA) synthase, and DHHA dehydrogenase (Fig. 1 and Supplementary Fig. 1). Chorismate is an ideal precursor for a new pathway because as a natural branch point for many primary and secondary metabolic processes it is already metabolically promiscuous and abundant in most cells^29, 30^. We therefore designed a third *de novo* biosynthesis pathway to NAD^+^ by combining the chorismate-to-3-HAA pathway with 3-HAA 3,4-dioxygenase (the enzyme converting 3-HAA to QA) and the common three-step process of pathways I and II converting QA to NAD^+^. This synthetic route was designated as C3N pathway based on its precursor <u>c</u>horismate, the key intermediate <u>3</u>-HAA and the final product <u>N</u>AD^+^, and the four enzymes converting chorismate to QA comprised the C3N module.

Stringent regulatory control of NAD^+^ *de novo* biosynthesis is directed mainly at the first biosynthetic genes like *nadB* and occurs at the transcriptional, translational, and post-translational levels^20–22^. By constructing the C3N pathway using defined promoters, ribosomal binding sites, and enzymes from secondary metabolism, these regulatory controls may theoretically be circumvented. Moreover, the recruitment of chorismate as the precursor for nicotinamide should decouple the C3N pathway from protein synthesis and make it suitable for use in engineered cells that overexpress proteins.

### Characterization of Pau20 as a DHHA dehydrogenase

Although the ADIC synthase and DHHA synthase enzymes catalyzing the first two steps of the C3N pathway have been well studied^31^, the third enzyme DHHA dehydrogenase has yet to be biochemically characterized (Supplementary Fig. 1). We chose the putative DHHA dehydrogenase Pau20 from the paulomycin biosynthetic gene cluster for characterization^27^. The *pau20* gene was inactivated by gene replacement in *Streptomyces paulus* NRRL 8115 to construct the *S. paulus pau20∷aac(3)IV* mutant (Supplementary Fig. 2a), in which the production of paulomycins was totally abolished. In feeding experiments of *S. paulus pau20∷aac(3)IV*, the production of paulomycins could be restored by 3-HAA but not by DHHA, indicating that Pau20 is responsible for the conversion of DHHA to 3-HAA (Fig. 2a). *N*-His_6_-tagged Pau20 was then overexpressed in *E. coli* BL21, purified, and incubated with DHHA and NAD^+^. Efficient oxidation of DHHA to 3-HAA was observed, verifying Pau20 as a DHHA dehydrogenase (Fig. 2b and 2c).

**Fig. 2.**
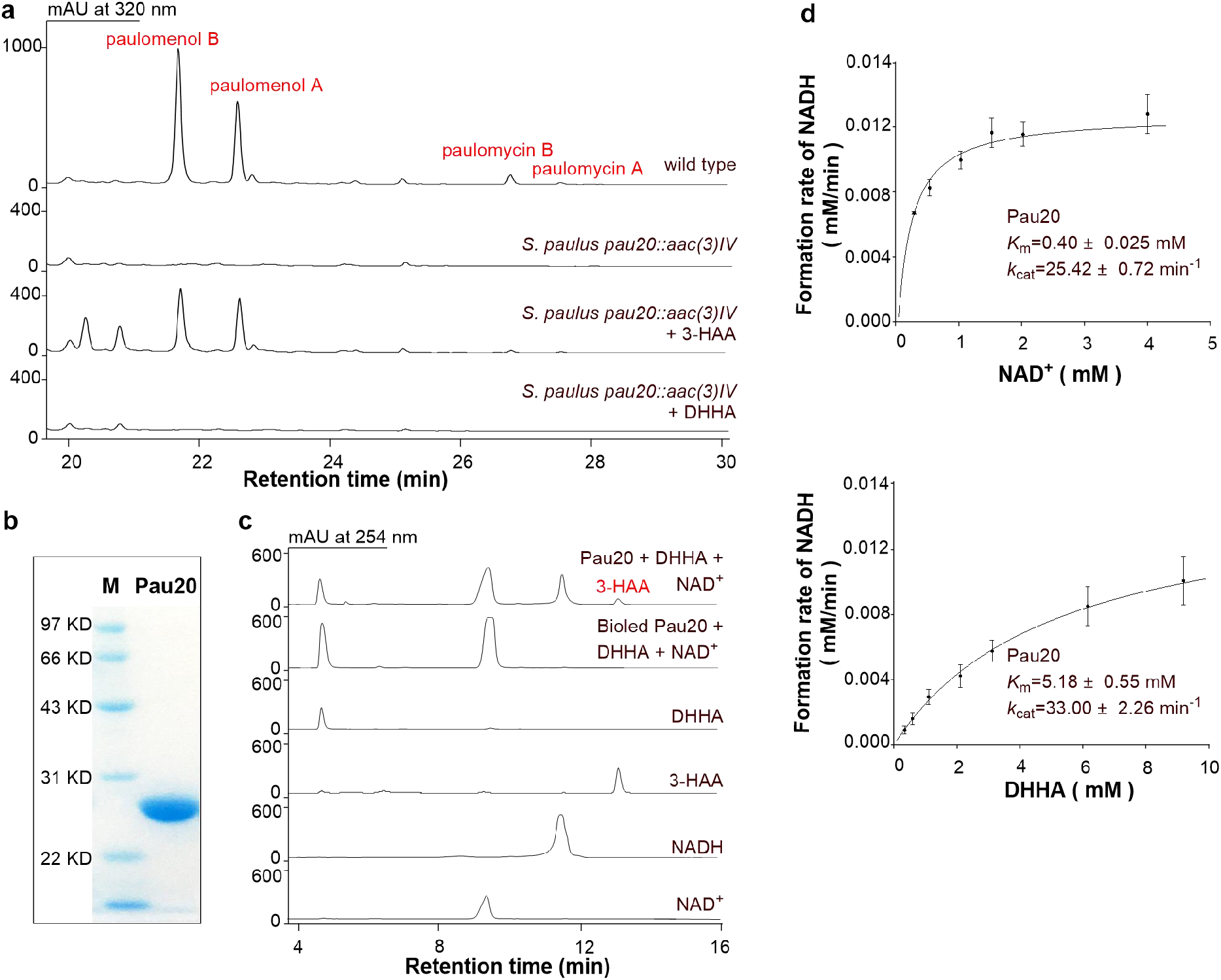
Characterization of Pau20 as a DHHA dehydrogenase. **a,** HPLC metabolite profiles of *S. paulus* wild type, the *pau20* inactivated mutant *S. paulus pau20∷aac(3)IV*, and *S. paulus pau20∷aac(3)IV* complemented with 3-HAA or DHHA. **b,** SDS-PAGE analysis of Pau20. M, protein marker; Pau20, *N*-His_6_-tagged Pau20. **c,** Representative enzymatic assays of Pau20. **d,** Steady-state kinetic analysis of Pau20.

Previous studies showed that 3-HAA is prone to oxidation under alkaline conditions and exposure to air^32^. Indeed, when the Pau20 assay was performed at buffers over pH 7.2, spontaneous oxidation of 3-HAA could be observed (Supplementary Fig. 3a). Therefore, Pau20 characterization was first performed at pH 7.0 and its optimal temperature of 37 °C (Supplementary Fig. 3b). Steady-state kinetic analysis of Pau20 under these conditions revealed Michaelis-Menten behavior for all substrates and kinetic constants consistent with Pau20 acting as an efficient DHHA dehydrogenase (Fig. 2d and Supplementary Fig. 3f). Characterization Pau20 *in vitro* not only supported a useful DHHA dehydrogenase for testing the designed C3N pathway, but for the first time unambiguously verified the chorismate to 3-HAA pathway originally proposed for a number of natural products.

### Constructing the C3N pathway in *E. coli*

The feasibility of the C3N pathway was initially tested by enabling 3-HAA production in *E. coli*, which lacks the kynurenine pathway and therefore cannot synthesize 3-HAA. A *pau20*-*phzDE* cassette consisting of *pau20* and two genes encoding the well-characterized ADIC synthase (PhzE) and DHHA synthase (PhzD) from the phenazine biosynthesis pathway of *Pseudomonas aeruginosa* PAO1^33, 34^ was synthesized and inserted into the medium-copy-number plasmid pXB1s downstream of the arabinose-inducible promoter *P_BAD_* to construct pXB1s-HAA. This plasmid was introduced into *E. coli* BW25113 to generate *E. coli* BW-pXB1s-HAA, which produced 3-HAA at a titer of 9.91 ± 0.14 mg/L (Fig. 3a).

**Fig. 3.**
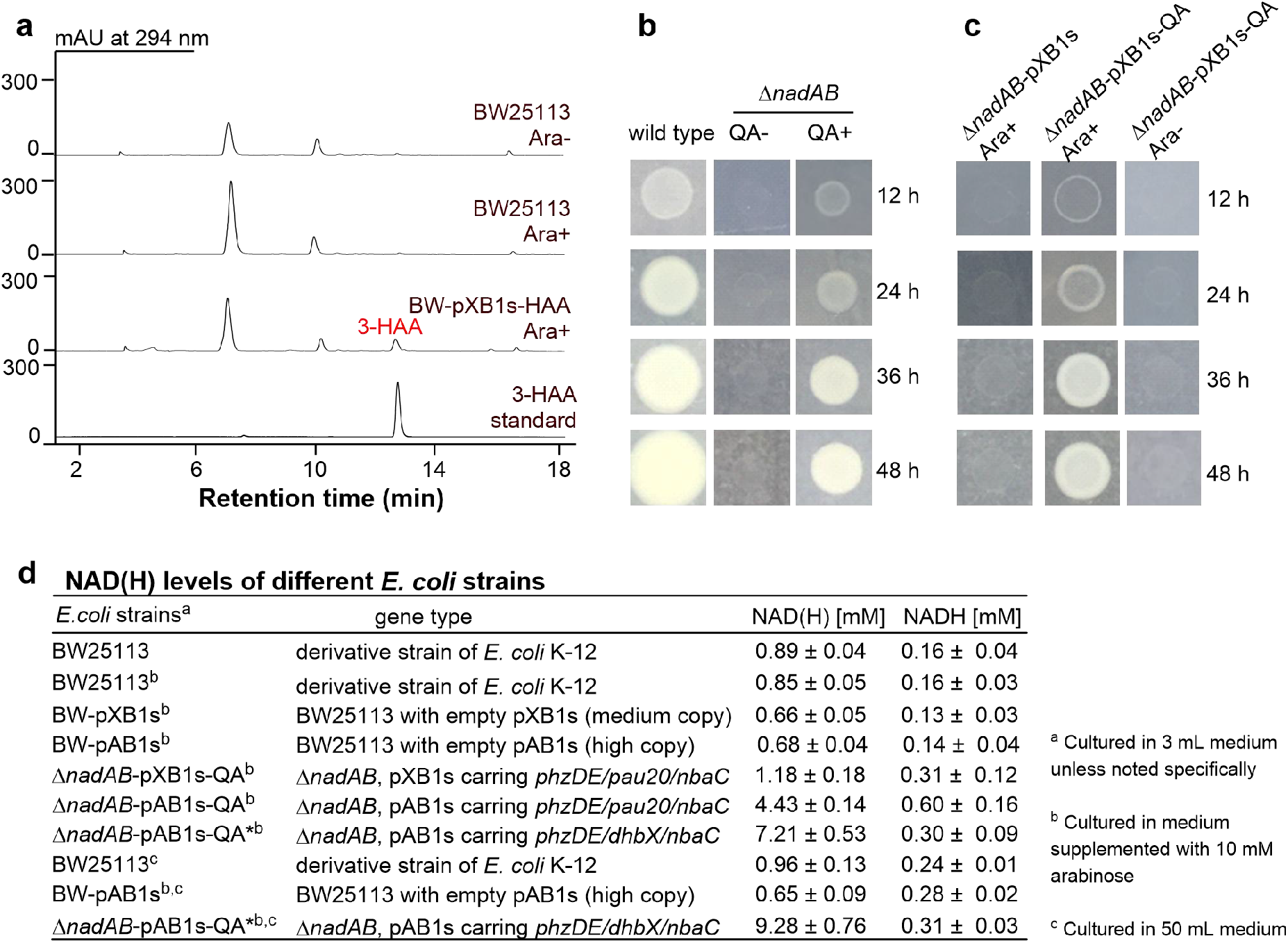
Construction of C3N pathway in *E. coli*. **a,** HPLC analysis of the metabolite profiles of *E. coli* BW25113 and BW-pXB1s-HAA with or without Ara (arabinose, 10mM). **b,** Growth of *E. coli* BW25113 (wild type) and Δ*nadAB* on M9 plates with or without QA (10 mM) addition. **c,** Growth of *E. coli ΔnadAB*-pXB1s-QA and the control strain *E. coli* Δ*nadAB*-pXB1s on M9 plates with or without arabinose (10 mM) addition. **d,** The cellular NAD(H) (NAD^+^ and NADH total) and NADH concentrations of different *E. coli* strains.

To establish the complete C3N pathway in *E. coli*, the *nbaC* gene encoding an efficient 3-HAA 3,4-dioxygenase from the aromatic compound degradation pathway of *Pseudomonas fluorescens* KU-7^35^ was inserted into plasmid pXB1s-HAA to generate pXB1s-QA, in which the four-gene cassette *nbaC-pau20*-*phzDE* dictates a C3N module converting chorismate to QA. In addition, to test whether cells could survive with the synthetic C3N pathway as the sole *de novo* source of NAD^+^, we constructed *E. coli* Δ*nadAB* (Supplementary Fig. 2b), a null mutant of NAD^+^ *de novo* biosynthesis, which was unable to grow on M9 plate unless QA was supplemented (Fig. 3b). Indeed, growth could be restored by introducing pXB1s-QA into *E. coli* Δ*nadAB* to afford *E. coli* Δ*nadAB*-pXB1s-QA, a recombinant strain with the complete C3N pathway. It grew well on M9 plates when the inducer arabinose was added. As a control, *E. coli* Δ*nadAB* with empty pXB1s could not grow on M9 plates even when supplemented with arabinose (Fig. 3c). Cellular NAD(H) levels measured at early stationary phase in liquid M9 medium with 10 mM arabinose were slightly higher in the C3N strain *E. coli* Δ*nadAB*-pXB1s-QA (1.18 ± 0.18 mM) than in *E. coli* BW25113 in M9 medium with (0.85 ± 0.05 mM) or without arabinose (0.89 ± 0.04 mM) (Fig. 3d). These results indicate that the C3N pathway was active in *E. coli* Δ*nadAB*-pXB1s-QA and could supply NAD^+^ efficiently for its growth.

### Optimizing the C3N pathway in *E. coli*

To assess the potential of the C3N pathway for improving cellular NAD(H) levels, we cloned the *nbaC-pau20*-*phzDE* cassette into the high-copy-number vector pAB1s. Transformation of the resultant plasmid into *E. coli* Δ*nadAB* afforded the strain *E. coli* Δ*nadAB*-pAB1s-QA with cellular NAD(H) concentrations as high as 4.43 ± 0.14 mM. HPLC analysis of the *E. coli* Δ*nadAB*-pAB1s-QA metabolic profile revealed that DHHA was accumulated in the fermentation broth (Supplementary Fig. 3c), implying that NAD(H) levels could be further increased by using more efficient DHHA dehydrogenases.

Six putative DHHA dehydrogenases (NatDB, CbxG, BomO, DhbX, CalB3, and StnN) were overproduced and purified as *N*-His_6_-tagged proteins and assayed along with Pau20 at pH 7.0 and 37 °C. All six enzymes displayed DHHA dehydrogenase activities, with DhbX, CalB3, and StnN outperforming the other three (Supplementary Fig. 3d-f). Steady-state kinetic analysis revealed that DhbX had the highest specificity constants among all tested DHHA dehydrogenases (Supplementary Fig. 3f). To exploit the efficient DhbX enzyme, we constructed plasmid pAB1s-QA* by synthesizing the *nbaC-dhbX*-*phzDE* cassette and cloning it into pAB1s. Introduction of pAB1s-QA* into *E. coli* Δ*nadAB* generated *E. coli* Δ*nadAB*-pAB1s-QA*, in which the accumulation of DHHA was clearly reduced (~35%) and its cellular NAD(H) concentration increased to 7.21 ± 0.53 mM (Fig. 3d and Supplementary Fig. 3c). When *E. coli* Δ*nadAB*-pAB1s-QA* was cultured in 250 mL flasks that contained 50 mL medium, the cellular NAD(H) concentration reached 9.28 ± 0.76 mM (Fig. 3d). Under the same cultivation conditions, the cellular NAD(H) levels of *E. coli* BW25113 and the control strain *E. coli* BW-pAB1s (*E. coli* BW25113 with empty pAB1s) were 0.96 ± 0.13 mM and 0.65 ± 0.09 mM, respectively.

The 9.7-fold increase in cellular NAD(H) concentration generated solely by the C3N pathway compares favorably with previous cofactor engineering efforts. Specifically, cellular NAD(H) levels could be enhanced 7-fold (to 7.03 mM) in *E. coli* BL21 by reinforcing NAD^+^ salvage via overexpression of the nicotinic acid phosphoribosyltransferase PncB and NAD^+^ synthetase NadE^15^. Other attempts to either limit NAD^+^ consumption or reinforce NAD^+^ *de novo* pathways achieved no more than 4-fold increases to cellular NAD(H) levels^17, 18^. Interestingly, an upper limit to cellular NAD(H) concentrations was proposed for the *E. coli* BW25113-derived strain YJE003. By inactivating *nadE* to block NAD^+^ synthesis and expressing the NAD(H) transporter *ntt4* gene to enable NAD^+^ intake, the maximum NAD(H) level of YJE003 was 9.6-fold (8.5 mM) higher than BW25113^36^. Similarly, our C3N-generated cellular NAD(H) concentration for *E. coli* Δ*nadAB*-pAB1s-QA* was 9.7-fold (9.3 mM) higher than BW25113, suggesting that the C3N pathway is optimized for NAD(H) yield and therefore harbors great potential for more general applications that require expanded cellular NAD(H) pools.

### Increasing DMP production *via* the C3N module

We envisioned that the C3N pathway converting chorismate to QA may represent a powerful tool to increase cellular NAD(H) levels. Specifically, we can consider the advantages of a packaged ‘C3N module’: (*i*) it can be easily introduced into targeted strains as a four-gene cassette; (*ii*) the cellular NAD(H) pools can be expanded significantly by the established C3N pathway in *E. coli*; and (*iii*) it is compatible with other NAD(H) level increasing strategies (Fig. 4a). We first tested this by introducing pAB1s-QA* into *E. coli* DMP, a cell factory for producing the food flavor additive 2,5-dimethylpyrazine (DMP), which is also the synthetic precursor of 5-methyl-2-pyrazinecarboxylic acid, a key component of widely used pharmaceuticals like Glipizide and Acipimox^37^. *E. coli* DMP overexpresses two genes encoding Tdh and SpNox. Tdh is an NAD^+^-dependent L-threonine-3-dehydrogenase that oxidizes L-threonine to (2*S*)-2-amino-3-oxobutanoate, which then spontaneously undergoes decarboxylation and dimerization to yield DMP^38^; SpNox is an NADH oxidase (H_2_O forming) for NAD^+^ recycling^39^. Compared with *E. coli* DMP-Con (*E. coli* DMP with empty pAB1s), the NAD(H) levels of the new strain *E. coli* C3N-DMP increased 2.7-fold, and DMP production was 3.8-fold higher (Fig. 4b). These results supported the application of the C3N module as a simple and effective tool to increase cellular NAD(H) levels and boost yields in microbial cell factories.

**Fig. 4.**
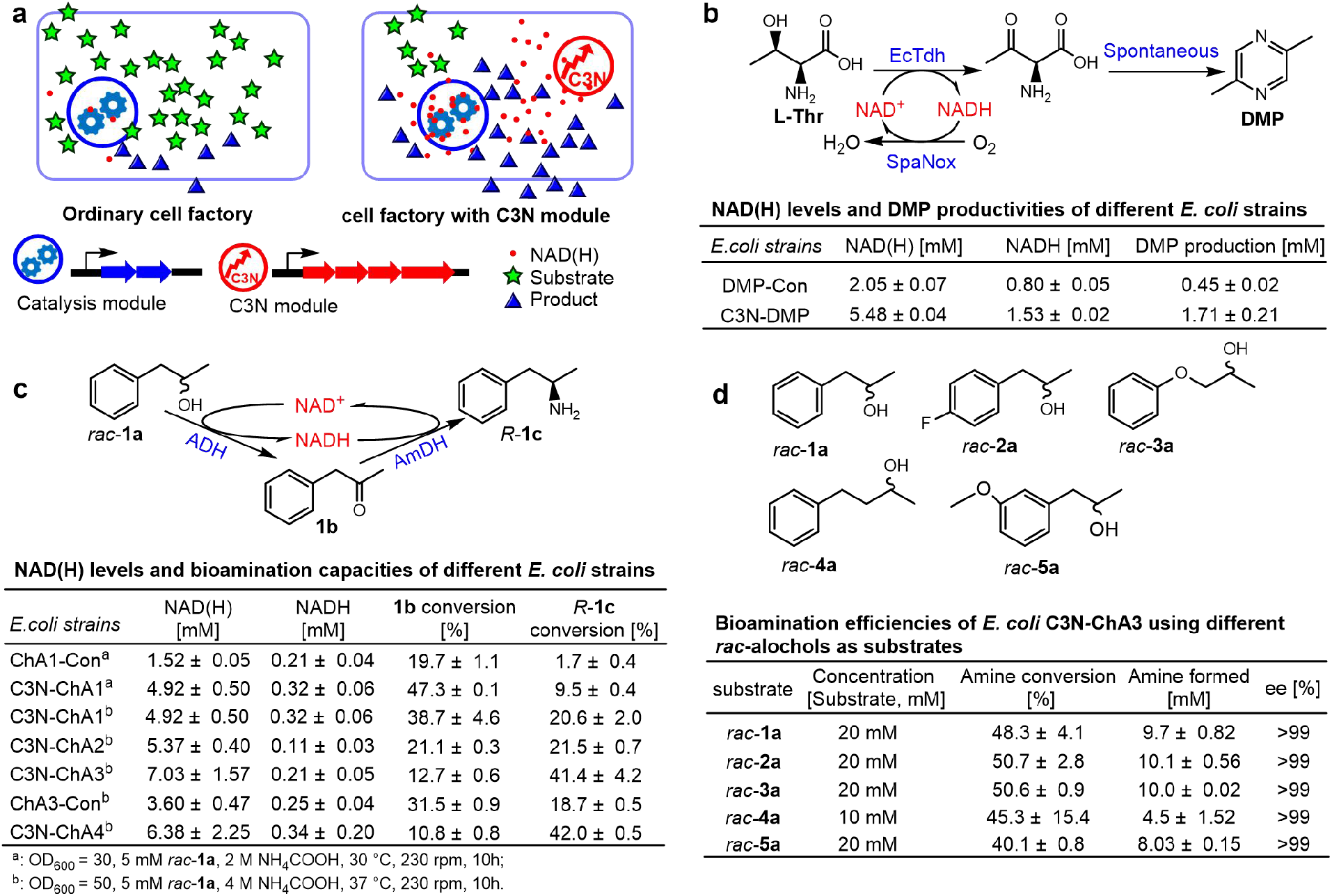
Improvement of the bioconversion efficiencies of cell factories by introducing the C3N module to increase cellular NAD(H) levels. **a,** General overview of improving the bioconversion efficiencies of cell factories by C3N module introduction. **b,** Improvement of DMP production by C3N module introduction. **c,** Evaluation of different C3N-based *E. coli* whole-cell bioamination systems. **d,** Evaluation of the bioamination efficiencies of *E. coli* C3N-ChA3 using different *rac*-alcohols as substrates. The bioconversions were performed in 100 mM KPi buffer (pH 8.5) with 10% DMSO, 4 M NH_4_COOH, OD_600_ of 50, at 37 °C, 230 rpm for 10 h.

### Construction of a C3N-based whole-cell system for preparing chiral amines

Encouraged by the convenience and efficacy of the C3N module towards boosting DMP production, we next sought to develop a C3N-based whole-cell system for preparing chiral amines. Chiral amines are important for synthetic intermediates of pharmaceuticals and other bioactive molecules^40^. Preparative scale amination of alcohols to enantiomerically pure chiral amines has been achieved using a biocatalytic hydrogen-borrowing cascade employing an NAD^+^-dependent alcohol dehydrogenase (ADH) coupled with an NADH-dependent amine dehydrogenase (AmDH) to enable efficient internal recycling of the nicotinamide coenzyme^40^ (Fig. 4c). A recent report described the first whole-cell conversion of alcohols to chiral amines by combining an enantioselective AA-ADH with a chimeric amine dehydrogenase Ch1-AmDH. The bioamination efficiencies in those whole-cell systems were enhanced by increasing cellular NAD(H) levels *via* the supplementation of NAD^+^ to lyophilized *E. coli* resting cells (external NAD^+^ cannot diffuse into cells without lyophilization) or by adding glucose to promote NADH recycling. This system could transform the starting alcohol (*S*)-**1a** to the chiral amine product (*R*)-**1c** (amphetamine) with 46 ± 14% yield (Fig. 4c). Adding the *R*-selective LBv-ADH enabled conversion of the racemic alcohol substrate *rac*-**1a** to product (*R*)-**1c** with 21 ± 6% yield^41^.

The critical role of the expensive reagent NAD(H) in this dual-enzyme system suggested it could benefit from C3N-boosted NAD(H) levels, and inspired us to develop an efficient whole-cell system for preparing chiral amines. We chose TeSADH, an ADH capable of accepting racemic secondary alcohols^42^, and CalAmDH, an AmDH with high activities toward phenylacetones^43^, to construct the dual-enzyme cascade. The medium copy number plasmid pCDF-*TesADH-CalAmDH* was transformed into *E. coli* BW25113(DE3) recombinant strains BW-C3N (harboring pAB1s-QA*) and BW-Con (harboring empty pAB1s) to afford *E. coli* C3N-ChA1 and the control strain ChA1-Con, respectively. The cellular NAD(H) concentration of *E. coli* C3N-ChA1 (4.92 ± 0.50 mM) was much higher than that of *E. coli* ChA1-Con (1.52 ± 0.03 mM) (Fig. 4c). With respect to increased production, the *rac*-**1a** substrate was turned over to product (*R*)-**1c** (*ee* > 99%) with 5.6-fold higher conversion for *E. coli* C3N-ChA1 (9.5 ± 0.4%) relative to *E. coli* ChA1-Con (1.7 ± 0.4%) (Fig. 4c).

Although incorporating the C3N module led to 5.6-fold higher conversion to the final product, further optimization was required to increase the bioamination efficiency of the whole-cell system. We therefore optimized the whole-cell bioconversion procedure for *E. coli* C3N-ChA1 with *rac*-**1a** (Supplementary Fig. 4a-d) and the selected optimal condition (100 mM KPi buffer, pH 8.5; 4 M NH_4_COOH; 5 mM *rac*-**1a**; OD_600_ = 50; 37 °C, 230 rpm for 10 h) doubled conversion to 20.6 ± 2.0 % of product (*R*)-**1c**. Further improvements were realized by additional gene copies to increase protein levels of the ADH-AmDH catalysis module. First, introduction of pACYC-*CalAmDH*, an expression plasmid of CalAmDH, into *E. coli* C3N-ChA1 afforded *E. coli* C3N-ChA2 with a similar conversion to (*R*)-**1c** of 21.5 ± 0.7 %. Next, the high copy number plasmid pRSF-*TesADH-CalAmDH* was transformed into *E. coli* BW-C3N to generate *E. coli* C3N-ChA3, which doubled conversion to 41.4 ± 4.2 % of product (*R*)-**1c**. Finally, *E. coli* C3N-ChA4 was obtained by transforming pACYC-*CalAmDH* into *E. coli* C3N-ChA3. It exhibited comparable bioamination capacity to *E. coli* C3N-ChA3 (Fig. 4c and Supplementary Fig. 4e).

Having constructed a set of strains with variable bioamination efficiencies, we sought to assess whether these efficiencies could be linked to C3N-generated NAD(H). Indeed, NAD(H) measurements revealed a clear positive correlation between bioamination efficiency and cellular NAD(H) levels of the resting cells (Fig. 4c). To further verify the effect of the C3N pathway on bioamination, a second control strain *E. coli* ChA3-Con was constructed by transforming pRSF-*TesADH-CalAmDH* into BW-Con. Under the optimized conditions the cellular NAD(H) levels of *E. coli* ChA3-Con (3.60 ± 0.47 mM) were much lower than that of *E. coli* C3N-ChA3 (7.03 ± 1.57 mM), and the conversion to final amine product was halved (18.7 ± 0.5%, (*R*)-**1c**) (Fig. 4c). When the concentration of *rac*-**1a** was increased to 20 mM, the conversion to product (*R*)-**1c** was much lower for *E. coli* ChA3-Con (6.7 ± 1.3%) than that for *E. coli* C3N-ChA3 (48.3 ± 4.1 %, Fig. 4d), further highlighting the important contribution of the C3N pathway.

We next explored the substrate promiscuity of the system using a series of phenyl-containing secondary alcohols (*rac*-**2a** to *rac*-**5a**). In all cases amination by *E. coli* C3N-ChA3 achieved remarkable amine conversions (40-50%) and optical purity (*ee* > 99% (*R*)), (Fig. 4d). To demonstrate the applicability of this whole-cell bioamination system at larger scale, preparative biotransformation of *rac*-**2a** (138 mg, 0.9 mmol) was carried out in a 30 mL mixture of *E. coli* C3N-ChA3 resting cells and resulted in 38.1 ± 1.9 % conversion to the product (*R*)-**2c**. When scaled up to 100 mL (using 462 mg *rac*-**2a**, 3 mmol), about 30% conversion was observed (producing 138 mg (*R*)-**2c**).

The C3N-based whole-cell system for chiral amine preparation revealed that the C3N module can be used as a powerful and expedient tool for cofactor engineering in *E. coli*. Moreover, we believe it has the potential to be used in a versatile range of host organisms because: (*i*) chorismate, the precursor of the C3N pathway, is a product of the shikimate pathway, which exists widely in bacteria, archaea, fungi, and plants^44^; (*ii*) the shared final three steps of NAD^+^ *de novo* biosynthesis pathways I and II exist in almost all organisms. In most cases, the only requirement for constructing the C3N pathway is to express the C3N module genes in the targeted organisms.

## Acknowledgments

This work was supported in part by the National Key R&D Program of China (2018YFA0901600 and 2018YFA0901900), the National Natural Science Foundation of China (31670032 and 31522001). Y.C. is an awardee for the ‘Hundred Talents Program’ of CAS.

## Author Contributions

Conceptualization, Y. D., X. L., and Y. C.; Methodology, Y. D., X. L., B. W., and Y. C.; Investigation, Y. D., X. L., P. L., M. W., J. L., Z. Z., and W. L.; Writing – Original Draft, Y. D., X. L., G. P. H., and Y. C.; Writing –Review & Editing, Y. D., G. P. H., B. W., Y. T., and Y. C.; Funding Acquisition, Y. C.; Resources, B. W., Y. T., and Y. C.; Supervision, Y. C.

## Methods

### Strains and plasmids

*E. coli* JM109 was used for DNA cloning. *E. coli* BL21(DE3) was used for expressing recombinant proteins. *E. coli* BW25113 and BW25113(DE3) were used for constructing C3N pathway and the C3N pathway-based whole-cell systems for preparing chiral amines, respectively. LB, M9 and M9Y media (M9 with 1 g/L yeast extract) were used for growth of *E. coli* strains at 37 °C except where noted specifically^45^. *S. paulus* NRRL 8115 and its mutants were grown on mannitol/soya (MS) agar for spore collections^46^. GS-7 medium was used as the seed culture and R5*α* medium was used for paulomycin production^47, 48^. All *Streptomyces* strains were cultured at 28 °C, 220 rpm. Strains and plasmids used in this study are summarized in the Supplementary Table 1.

### General DNA manipulations and sequence analyses

DNA synthesis and sequencing were performed in Generay Biotech (Shanghai, China) and Biosune (Shanghai, China) respectively. PCRs were performed with KOD-Plus DNA polymerase (Toyobo, Osaka, Japan) or Q5 DNA polymerase (NEB, Beijing). The one-step cloning method was employed in plasmid construction according to the manufacturer’s instructions (Vazyme, Nanjing, China). Restriction enzyme digestions, ligations and transformations were carried out following general methods^45^. *E. coli-Streptomyces* conjugations were performed as described^46^. A BLASTP search was used to find protein homologs (https://blast.ncbi.nlm.nih.gov/Blast.cgi). Protein sequence alignments were performed using Clustal Omega (https://www.ebi.ac.uk/Tools/msa/clustalo/).

### Construction of *S. paulus pau20∷aac(3)IV*

A blue-white screening-based gene inactivation system was used to inactivate gene *pau20*^49^. The 1.9-kb upstream fragment and the 2.2-kb downstream fragment of *pau20* were amplified using primer pairs pau20-s2/pau18R and pau22-R/pau20-R and inserted into the *Mun*I and *Bln*I sites of pCIMt002 respectively to afford the *pau20* disruption plasmid, which was then introduced into *S. paulus* NRRL 8115 *via E. coli-Streptomyces* conjugation. Both blue and white exconjugants were obtained on the plate. One of the white exconjugants with apramycin resistance was selected as the desired *pau20* gene inactivation mutants *S. paulus pau20∷aac(3)IV*, which was verified by PCR with primer pair pau20-s2/ pau20-R (Supplementary Fig. 2a).

### Production of paulomycins

For paulomycin production, 50 *μ*L spores of *S. paulus* NRRL 8115 or *pau20∷aac(3)IV* were inoculated into GS-7 liquid medium and cultured at 28 °C, 220 rpm for 2 days. The resulting seed culture was inoculated into 50 mL R5α liquid medium at a 2% ratio (v/v). After 4 days fermentation, the broth was harvested by centrifugation, extracted with 50 mL ethyl acetate three times, and concentrated *in vacuo*. The samples were re-dissolved in 1 mL acetonitrile and subjected to HPLC analysis.

### Complementation of *S. paulus pau20∷aac(3)IV* by feeding DHHA or 3-HAA

For the feeding experiments, *S. paulus pau20∷aac(3)IV* was cultured at the paulomycin production condition. DHHA or 3-HAA was added to the cultures to a final concentration of 2.5 mM two days after seed inoculation. After culturing at 28 °C, 220 rpm for 2 days, the broth was harvested, extracted, and concentrated as described above. The samples were re-dissolved in 1 mL acetonitrile and analyzed by HPLC.

### Expression and purification of the DHHA dehydrogenases

The 0.7-kb *pau20* gene was cloned using the *S. paulus* NRRL 8115 genome as a template with primer pair pauN10ES/pauN10ER, verified by sequencing, and inserted into the *Nde*I*/Bam*HI sites of pET-28a to generate pET-28a-*pau20*. A single transformant of *E. coli* BL21(DE3)/pET-28a-*pau20* was inoculated into LB medium with 50 *μ*g/mL kanamycin and cultured overnight at 37 °C, 220 rpm. The overnight seed culture was inoculated to the same medium at 1% ratio (v/v) and incubated at 37 °C, 220 rpm until OD_600_ reached 0.6. Expression of Pau20 was then induced by the addition of isopropyl-*β*-thiogalactoside (IPTG, 0.05 mM) and further cultured at 16 °C, 180 rpm for 12-16 h.

Purification of Pau20 was carried out with the Ni-NTA affinity column at 4 °C following the manufacturer’s instructions. The *E. coli* cells harvested by centrifugation (4 °C, 3000 × g, 10 min) were re-suspended in lysis buffer (20 mM Tris-HCl, 500 mM NaCl, 5 mM imidazole, pH 7.9) and burst by ultrasonication. After removing the cell debris by centrifugation (4 °C, 16,000 × g, 30 min), the remaining supernatant was subjected to the Ni-NTA affinity column and washed with washing buffer (lysis buffer with 60 mM imidazole) and then elution buffer (lysis buffer with 500 mM imidazole). The purified protein was desalted and concentrated by ultrafiltration using a Millipore centrifugal filter (15 mL, 10 kDa MWCO) and stored at −80 °C in phosphate-buffered saline (PBS) buffer (pH 7.4) with 20% glycerol.

The other six DHHA dehydrogenase genes, *natDB* (NatDB Accession: CEK42820), *cbxG* (CbxG Accession: KDQ70111), *dhbX* (DhbX Accession: CDG76955), *bomO* (BomO Accession: ALE27507), *calB3* (ClaB3 Accession: AEH42481), and *stnN* (StnN Accession: AFW04566), were synthesized and inserted into the *Nde*I*/Bam*HI sites of pET-28a to afford plasmids pET-28a-*natDB*, pET-28a-*cbxG*, pET-28a-*dhbX*, pET-28a-*bomO*, pET-28a-*calB3*, and pET-28a-*stnN* respectively. The plasmids were transformed into *E. coli* BL21(DE3) individually to afford the protein expression strains. Protein expression and purification of the six DHHA dehydrogenases were carried out using the same procedures as those of Pau20. The purified enzymes were stored at −80 °C in PBS buffer with 20% glycerol. Protein concentrations were measured with a BCA protein assay kit (Takara, Dalian, China).

### Enzymatic assays of the DHHA dehydrogenases

The DHHA dehydrogenase activity of Pau20 was tested in a 100 *μ*L mixture containing 50 mM PBS buffer (pH 7.4), 2 mM NAD^+^, 2 mM DHHA, and 5 *μ*M Pau20 at 30 °C for 3 h. The reaction was quenched by equal volume of CHCl_3_. After centrifugation, the aqueous phase of the reaction (20 *μ*L) was analyzed by HPLC. The pH value optimization of Pau20 was carried out at 37 °C in buffers of pH ranging from 6.6 to 8.5 (200 mM phosphate buffer for pH 6.0-7.6, and 50 mM Tris-HCl buffer for pH 8.5) for 30 min. Temperature optimization was performed in a 100 *μ*L reaction mixture in 200 mM phosphate buffer (pH 7.0) for 30 min. The steady-state kinetic analysis of Pau20 was carried out in 200 mM phosphate buffer (pH 7.0) at 37 °C. The enzymatic assays of the other six DHHA dehydrogenases and the steady-state kinetic analyses of DhbX, CalB3, and StnN were all carried out using the same reaction conditions as Pau20 in 200 mM phosphate buffer at 37 °C.

### Construction and evaluation of *E. coli* BW-pXB1s-HAA

A 2.6-kb DNA fragment containing the 0.6-kb *phzD* (PhzD Accession: AAC64487) and the 1.9-kb *phzE* genes (PhzE Accession: AAC64488) was synthesized and the homolog sequences for ligation independent cloning at its both ends were added by a PCR amplification using primer pair pHAAphzDE-F/pHAAphzDE-R. The 0.7-kb DNA fragment containing the *pau20* gene was amplified from the *S. paulus* NRRL 8115 genome using primer pair pHAApau20-F/pHAApau20-R. The two DNA fragments were inserted into the *Nco*I*/Eco*RI sites of pXB1s (p15A origin, medium-copy-number) using one-step cloning kit (Vazyme, Nanjing, China) to generate pXB1s-HAA. *E. coli* BW-pXB1s-HAA was constructed by introducing pXB1s-HAA into *E. coli* BW25113. *E. coli* BW-pXB1s-HAA was cultured in 3 mL LB medium with 50 *μ*g/mL streptomycin at 37 °C overnight. The seed culture was then inoculated into 50 mL M9 medium with 50 *μ*g/mL streptomycin at 1% ratio (v/v). After cultured at 37 °C, 220 rpm for 3 h, 10 mM arabinose was added and the culture was further fermented at 37°C, 220 rpm for 20 h. The broth was centrifuged to remove the cell pellet and the supernatant was subjected to HPLC analysis directly.

### Construction of *E. coli* Δ*nadAB*-pXB1s-QA, Δ*nadAB*-pAB1s-QA and Δ*nadAB*-pAB1s-QA*

*E. coli* BW25113 *nadA∷aph* is a commercially available *nadA* disrupted mutant, in which gene *nadA* is replaced by the kanamycin resistance gene cassette *aph* flanked by two FRT (FLP recognition targets) sites. After introducing plasmid pCP20 with the FLP-recombinase gene into *E. coli* BW25113 *nadA∷aph*, one of the kanamycin sensitive strains was selected as the *nadA* deleted mutant *E. coli* Δ*nadA* and was verified by PCR using primer pair nadA-F/nadA-R. To knock-out *nadAB*, P1 bacteriophage was used for dissociation of the *nadB* gene inactivated mutant *E. coli* BW25113 *nadB∷aph* and then introduced into *E. coli* Δ*nadA* by transduction (Thomason et al., 2007). One of the kanamycin resistant strains was selected and verified as *E. coli* Δ*nadAB* by PCR analysis with primer pairs nadA-F/nadA-R and nadB-F/nadB-R (Supplementary Fig.2b). The 0.6-kb *nbaC* (NbaC Accession: WP_006094629) was synthesized and the homolog sequences for ligation independent cloning at its both ends were added by a PCR amplification using primer pair NbaC-F/NbaC-R. The *nbaC* gene was then inserted into the *Nco*I site of pXB1s-HAA to generate pXB1s-QA, which was transformed into *E. coli* Δ*nadAB* to obtain *E. coli* Δ*nadAB*-pXB1s-QA.

Plasmids pAB1s-HAA (ColE1 origin, high-copy-number) was constructed similarly as that of pXB1s-HAA by insertion of the 2.6-kb fragment containing genes *phzDE* and the 0.7-kb fragment containing gene *pau20* into the *Nco*I*/Eco*RI sites of pAB1s. The 0.6-kb *nbaC* gene was then inserted into the *Nco*I sites of pAB1s-HAA to generate pAB1s-QA, which was transformed into *E. coli* Δ*nadAB* to obtain *E. coli* Δ*nadAB*-pAB1s-QA.

The 0.7-kb *dhbX* gene was synthesized and the homolog sequences for ligation independent cloning at its both ends were added by a PCR amplification using primer pair DhbX-F/DhbX-R. It was then inserted into the *Xba*I site of pAB1s-QA to replace the *pau20* gene (excised by *Xba*I digestion) to generate pAB1s-QA*, which was transformed into *E. coli* Δ*nadAB* to obtain *E. coli* Δ*nadAB*-pAB1s-QA*.

### Growth of *E. coli* strains on M9 plates

The *E. coli* strains were cultured in 3 mL LB medium with appropriate antibiotics at 37 °C, 220 rpm overnight and harvested by centrifugation (4 °C, 3000 × g, 5 min). To remove extracellular NAD(H) and its salvage pathway precursors, the cells were washed twice with ddH_2_O and diluted to an OD_600_ of 0.1. For the growth experiments on agar plates, 2 *μ*L dilutions (OD_600_ of 0.1) of different *E. coli* strains were spotted onto M9 agar with varied supplements. *E. coli* Δ*nadAB* was spotted onto M9 agar supplemented with or without 10 mM QA; *E. coli* Δ*nadAB*-pXB1s-QA and Δ*nadAB*-pAB1s-QA and their controls with empty vectors were inoculated onto M9 agar (50 *μ*g/mL streptomycin) with or without 10 mM arabinose. The *E. coli* strains were grown at 37 °C.

### Construction and evaluation of the DMP cell factory

A 2.4-kb DNA fragment containing the 1.0-kb *EcTdh* (EcTDH, Accession No.: WP_000646007) and the 1.4-kb *SpaNox* (SpaNox, Accession No.: WP_002989788) genes was synthesized and inserted into the *Nco*I*/Xho*I sites of pRSFDuet-1 to afford plasmid pRSF-*EcTdh*-*SpaNox*. *E. coli* C3N-DMP and the control strain *E. coli* DMP-Con were constructed by introducing pRSF-*EcTdh*-*SpaNox* into *E. coli* BL21-C3N (containing plasmid pAB1s-QA*) and BL21-Con (containing plasmid pAB1s), respectively. Briefly, *E. coli* strains C3N-DMP and DMP-Con were cultured in 3 mL LB medium with 50 *μ*g/mL streptomycin and 50 *μ*g/mL kanamycin at 37 °C overnight. The seed cultures were inoculated into 100 mL M9Y medium (M9 Medium containing 0.1% yeast extract) with 50 *μ*g/mL streptomycin and 50 *μ*g/mL kanamycin at 1% ratio (v/v). After culturing at 37 °C, 220 rpm for 3 h, 0.1 mM IPTG and 10 mM arabinose were added and the cultures were further cultured at 37 °C, 220 rpm for 24 h. The cells were then collected by centrifugation (4 °C, 3,000 × g, 20 min), washed with ddH_2_O and resuspend in 100 mM Tris-HCl buffer (pH 8.0). The whole-cell conversions were started in 2 mL Eppendorf tubes containing 1 mL 100 mM Tris-HCl buffer (pH 8.0) under 28 °C and 230 rpm for 12 h. The initial concentrations of L-threonine and the resting cells were set as 200 mM and OD_600_ of 35, respectively. To detect the production of DMP, the reaction samples were centrifuged (4 °C, 16,000 × g, 10 min) to remove the pellets, diluted 4 times with acetonitrile, and subjected to HPLC analysis.

### Construction of *E. coli* strains BW-C3N-ChA1 to BW-C3N-ChA4

The codon-optimized alcohol dehydrogenase *TesADH* gene (TesADH, Accession No.: WP_041589967, point mutations W110A and G198D were modified according to Thompson et al., 2017) and amine dehydrogenase *CalAmDH* gene (CalAmDH, Accession No.: WP_007505854, point mutations K68S and N266L were modified according to previous reported^43^) were synthesized and inserted into the *Nco*I*/Bam*HI sites of pET-28a to afford plasmids pET28a-*TesADH* and pET28a-*CalAmDH*, respectively. The 1.1-kb DNA fragment containing the *TesADH* gene was amplified from the pET28a-*TesADH* using primers T7/Cal-P23R and inserted into the *Nco*I/*Sal*I site of pCDFduet-1a (derivate from pCDFduet-1, Amp^R^) to generate pCDF-*TesADH*. The 1.1-kb DNA fragment containing the *CalAmDH* gene was amplified from the pET28a-*CalAmDH* using primers CALRS-F/CALRS-R and inserted into the *Nde*I/*Xho*I site of pCDF-*TesADH* to generate pCDF-*TesADH-CalAmDH*. *E. coli* BW-C3N-ChA1 was obtained by introducing pCDF-*TesADH-CalAmDH* into *E. coli* BW-C3N (*E. coli* BW25113(DE3) containing plasmid pAB1s-QA*), and the control strain *E. coli* BW-ChA1-Con was constructed by introducing the same plasmid into *E. coli* BW-Con (*E. coli* BW25113(DE3) containing plasmid pAB1s). To construct *E. coli* BW-C3N-ChA2, the *CalAmDH* gene was cloned from pET28a-*CalAmDH* with primers T7/Cal-his-R and inserted into the *Nco*I/*Sal*I site of pACYduet-1 to generate pACYC-*CalAmDH*, and the resultant plasmid was introduced into *E. coli* BW-C3N-ChA1. The 2.3-kb DNA fragment containing genes *TesADH* and *CalAmDH* was amplified from pCDF-*TesADH-CalAmDH* using primers T7/Cal-P23R and cloned into the *Nco*I/*Xho*I site of pRSFduet-1 (RSF origin, high-copy-number) to generate pRSF-*TesADH-CalAmDH*, which was transformed into *E. coli* BW-C3N and BW-Con respectively to afford *E. coli* BW-C3N-ChA3 and the control strain BW-ChA3-Con. *E. coli* BW-C3N-ChA4 was obtained by introducing pACYC-*CalAmDH* into *E. coli* BW-C3N-ChA3.

### General procedures for bioamination of *rac*-alcohols

The engineered *E. coli* strains was inoculated in 3 mL LB broth with appropriate antibiotics when necessary (50 *μ*g/mL kanamycin, 50 *μ*g/mL streptomycin, 100 *μ*g/mL ampicillin, 20 *μ*g/mL chloramphenicol) and cultured at 37 °C, 220 rpm overnight. The seed cultures were then inoculated into 100 mL M9Y medium at 1% ratio (v/v). After culturing at 37 °C, 220 rpm for 3 h, 0.1 mM IPTG and 10 mM arabinose were added and the cultures were further fermented for 24 h. The cells were then harvested by centrifugation (4 °C, 3,000 × g, 20 min), washed by ddH_2_O and re-suspended in appropriate buffers for the whole-cell bioconversions. For comparison of the bioamination efficiencies of *E. coli* BW-ChA1-Con and BW-C3N-ChA1, the bioconversions were carried out in 100 mM KPi buffer (pH 8.5) with 10% DMSO in 1 mL reaction volume with 5 mM **1a**, 2 M NH_4_COOH under 30 °C, 230 rpm for 10 h. To find an optimal bioamination procedure, varied buffers (100 mM KPi, 100 mM Tris-HCl, and 2 M NH_3_/NH_4_Cl (all buffers were adjusted to pH 8.5 and with 10% DMSO), concentrations of NH_4_^+^ (range from 0.5 M to 4 M), amounts of resting cells (OD_600_ of 30, 50 and 70), and reaction temperatures (25 °C, 30 °C, 37 °C, and 42 °C) were sequentially optimized in 1 mL reactions with 5 mM substrate **1a** and keeping the agitation at 230 rpm.

The analytical scale bioamination reactions of *E. coli* BW-C3N-ChA1 to BW-C3N-ChA4 with different alcohol substrates were then carried out under the optimal condition (100 mM KPi buffer (pH 8.5) with 10% DMSO, 5 or 20 mM *rac*-alcohols, 4 M NH_4_COOH, OD_600_ of 50, 37 °C, 230 rpm) in 2 mL Eppendorf tubes containing 1 mL reaction volume for 10 h. To scale up the biotransformation system, *E. coli* BW-NAD-ChA3 was cultured, collected and resuspended in a 250 mL flask containing 30 mL or 100 mL KPi buffer (100 mM, pH 8.5) with 10% DMSO, 30 mM *rac*-**2a**, 4 M NH_4_COOH, OD_600_ of 50 resting cells, and the whole-cell bioconversions were performed at 37 °C, 230 rpm for 24 h. All biotransformation reactions were quenched using 100 *μ*L 10 M KOH per mL reaction mixture. After centrifugation at 16,000 × g for 10 min, 200 *μ*L supernatant was extracted by the same volume of ethyl acetate three times. The organic phase (20 *μ*L) was then subjected to HPLC analysis.

### Derivatization of the chiral amine samples

The enantiomeric excess of the amine products was analyzed using the FDAA (Marfey’s Reagent) pre-column derivatization method. Briefly, the reaction mixture (200 *μ*L) was extracted by ethyl acetate, dried in vacuum, and re-suspended in 20 *μ*L acetone. To perform the derivatization, the sample was incubated with 40 *μ*L FDAA (1% pre-dissolved in acetone, w/v) and 40 *μ*L 1 M NaHCO_3_ at 40 °C for 1 h. The reaction was quenched by adding 80 *μ*L 1 M HCl and then diluted with 800 *μ*L ethyl acetate. The supernatant was subjected to HPLC analysis after centrifugation.

### Measurement of intracellular NAD(H) concentrations

The *E. coli* strain was cultivated in 3 mL LB medium with appropriate antibiotics at 37 °C, 220 rpm overnight, and the overnight culture was then inoculated into 3 mL or 50 mL M9 medium at 1% ratio (v/v) and grown at 37°C, 220 rpm for 18-24 h. To measure the intracellular NAD(H) concentration, the cells were harvested by centrifugation (4 °C, 3,000 × g, 5 min) at the time point of four hours after entering stationary phase. The cell pellets were washed with ice-cold PBS buffer, re-suspended in the same buffer and adjusted to OD_600_ of 1.0. The concentrations of NAD(H) (total NAD^+^/NADH) and NADH only were measured by the NAD^+^/NADH assay kit (Abcam, Cambridge, UK) according to the manufacturer’s instructions. *E. coli* cell concentration was calculated as 10^9^ cells/mL at OD_600_ of 1.0 and the volume of one *E. coli* cell was 10^−12^ mL^36, 50^ (Zhou et al., 2011; Brumaghim et al., 2003).

### Spectroscopic analysis

HPLC analysis was carried out on a Shimadzu HPLC system (Shimadzu, Kyoto, Japan). The analysis of paulomycin production was performed using an Apollo C18 column (5 *μ*m, 4.6 × 250 mm, Alltech, Deerfield, IL, USA), which was developed with a linear gradient using acetonitrile and water with 0.1% trifluoroacetic acid at a flow rate of 0.8 mL/min. The ratio of acetonitrile was maintained at 5% for 5 min and changed linearly from 5% to 90% over 5-25 min and from 90% to 100% over 25-30 min. The detection wavelength was 320 nm.

The formation of NADH in the DHHA dehydrogenase reactions was monitored by BioTek Synergy H4 Hybrid Reader (BioTek, Vermont, USA) at 340 nm. HPLC analysis of the DHHA dehydrogenase reactions and production of 3-HAA in *E. coli* was carried out with a ZORBAX SB-Aq StableBond Analytical column (5 *μ*m, 4.6 × 250 mm, Agilent Teologies, Santa Clara, CA, USA). A linear gradient of acetonitrile and water with 0.1% trifluoroacetic acid was used for development of the column at a flow rate of 0.8 mL/min. The ratio of acetonitrile was changed linearly from 1% to 50% over 0-30 min, from 50% to 100% over 30-32 min, and maintained at 100% for 5 min. The detection wavelength was 254 nm or 294 nm.

The production of DMP was analyzed using an Apollo C18 column (5 *μ*m, 4.6 × 250 mm, Alltech, Deerfield, IL, USA) using an isocratic elution of acetonitrile/water (0.1% trifluoroacetate, v/v) (20:80, v/v) at 0.8 mL/min. The detection wavelength was 275 nm.

The bioamination samples were analyzed using an Apollo C18 column (5 *μ*m, 4.6 mm × 250 mm, Alltech, Deerfield, IL, USA) using an isocratic elution of acetonitrile/water (0.1% trifluoroacetate, v/v) (30:70, v/v) at 0.8 mL/min. The detection wavelength was 254 nm. HPLC analysis of the enantiomeric excess of chiral amine products was carried out with the same Apollo C18 column with a linear gradient using acetonitrile with 0.1% formic acid and water with 0.1% formic acid at a flow rate of 1 mL/min. The percentage of acetonitrile started being held constant at 20% for 10 min, changed from 20 to 60% over 30 min and from 60 to 100% over 5 min. The detection wavelength was 340 nm.

LC-MS analysis was performed with the ZORBAX SB-Aq StableBond Analytical column (5 *μ*m, 4.6 × 250 mm, Agilent Teologies, Santa Clara, CA, USA) on an Agilent 1260/6460 Triple-Quadrupole LC/MS system (Santa Clara, CA, USA).

### Chemicals

The racemic alcohols **3a** and **5a** were chemically synthesized by reduction of the commercially available ketones **3b** and **5b**; the ketone **1b** was chemically synthesized by oxidation of **1a**. Enantiomeric standard (*S*) and (*R*) amines of **1c-5c** were synthesized by stereoselective amination using previously described enzymatic methods^51^.

## References

1. Begley, T. P., Kinsland, C., Mehl, R. A., Osterman, A. & Dorrestein, P. The biosynthesis of nicotinamide adenine dinucleotides in bacteria. Vitam. Horm. 61, 103–119 (2001).

2. Kurnasov, O. et al. NAD biosynthesis: Identification of the tryptophan to quinolinate pathway in bacteria. Chem. Biol. 10, 1195–1204 (2003).

3. Rongvaux, A., Andris, F., Van Gool, F. & Leo, O. Reconstructing eukaryotic NAD metabolism. Bioessays 25, 683–690 (2003).

4. Bieganowski, P. & Brenner, C. Discoveries of nicotinamide riboside as a nutrient and conserved NRK genes establish a Preiss-Handler independent route to NAD^+^ in fungi and humans. Cell 117, 495–502 (2004).

5. Vidal, L. S., Kelly, C. L., Mordaka, P. M. & Heap, J. T. Review of NAD(P)H-dependent oxidoreductases: properties, engineering and application. Biochim. Biophys. Acta. Proteins Proteom 1866, 327–347 (2018).

6. Wang, Y. P., San, K. Y. & Bennett, G. N. Cofactor engineering for advancing chemical biotechnology. Curr. Opin. Biotech. 24, 994–999 (2013).

7. Wang, M., Chen, B., Fang, Y. & Tan, T. Cofactor engineering for more efficient production of chemicals and biofuels. Biotechnol. Adv. 35, 1032–1039 (2017).

8. Belas, R. et al. Bacterial bioluminescence: isolation and expression of the luciferase genes from *Vibrio harveyi*. Science 218, 791–793 (1982).

9. Shah, G. M. et al. Biochemical assessment of niacin deficiency among carcinoid cancer patients. Am. J. Gastroenterol. 100, 2307–2314 (2005).

10. Berger, F., Ramirez-Hernandez, M. H. & Ziegler, M. The new life of a centenarian: signalling functions of NAD(P). Trends Biochem. Sci. 29, 111–118 (2004).

11. Bentle, M. S., Reinicke, K. E., Bey, E. A., Spitz, D. R. & Boothman, D. A. Calcium-dependent modulation of poly(ADP-ribose) polymerase-1 alters cellular metabolism and DNA repair. J. Biol. Chem. 281, 33684–33696 (2006).

12. Imai, S. & Guarente, L. Ten years of NAD-dependent SIR2 family deacetylases: implications for metabolic diseases. Trends Pharmacol. Sci. 31, 212–220 (2010).

13. Zhang, H. B. et al. NAD^+^ repletion improves mitochondrial and stem cell function and enhances life span in mice. Science 352, 1436–1443 (2016).

14. Katsyuba, E. et al. *De novo* NAD^+^ synthesis enhances mitochondrial function and improves health. Nature 563, 354 (2018).

15. Heuser, F., Schroer, K., Lutz, S., Bringer-Meyer, S. & Sahm, H. Enhancement of the NAD(P)(H) pool in *Escherichia coli* for biotransformation. Eng. Life Sci. 7, 343–353 (2007).

16. Zhou, Y. et al. Engineering NAD^+^ availability for *Escherichia coli* whole-cell biocatalysis: a case study for dihydroxyacetone production. Microb. Cell Fact. 12(2013).

17. Han, Q. & Eiteman, M. A. Enhancement of NAD(H) pool for formation of oxidized biochemicals in *Escherichia coli*. J. Ind. Microbiol. Biot. 45, 939–950 (2018).

18. Liao, Z., Yang, X., Fu, H. & Wang, J. The significance of aspartate on NAD(H) biosynthesis and ABE fermentation in *Clostridium acetobutylicum* ATCC 824. AMB Express 9 (2019).

19. Li, F. et al. Modular engineering to increase intracellular NAD(H/^+^) promotes rate of extracellular electron transfer of *Shewanella oneidensis*. Nat. Commun. 9 (2018).

20. Rodionov, D. A. et al. Transcriptional regulation of NAD metabolism in bacteria: NrtR family of Nudix-related regulators. Nucleic Acids Res. 36, 2047–2059 (2008).

21. Malkowski, S. N., Spencer, T. C. & Breaker, R. R. Evidence that the *nadA* motif is a bacterial riboswitch for the ubiquitous enzyme cofactor NAD^+^. RNA 25, 1616–1627 (2019).

22. Nasu, S., Wicks, F. D. & Gholson, R. K. L-aspartate oxidase, a newly discovered enzyme of *Escherichia coli*, is the b protein of quinolinate synthetase. J. Biol. Chem. 257, 626–632 (1982).

23. Wu, Q. L. et al. Characterization of the biosynthesis gene cluster for the pyrrole polyether antibiotic calcimycin (A23187) in *Streptomyces chartreusis* NRRL 3882. Antimicrob. Agents. Chemother. 55, 974–982 (2011).

24. Xu, F. et al. Characterization of streptonigrin biosynthesis reveals a cryptic carboxyl methylation and an unusual oxidative cleavage of a N-C bond. J. Am. Chem. Soc. 135, 1739–1748 (2013).

25. Schneditz, G. et al. Enterotoxicity of a nonribosomal peptide causes antibiotic-associated colitis. Proc. Natl. Acad. Sci. USA 111, 13181–13186 (2014).

26. Cano-Prieto, C. et al. Genome mining of *Streptomyces* sp. tü 6176: characterization of the nataxazole biosynthesis pathway. Chembiochem 16, 1461–1473 (2015).

27. Li, J., Xie, Z., Wang, M., Ai, G. & Chen, Y. Identification and analysis of the paulomycin biosynthetic gene cluster and titer improvement of the paulomycins in *Streptomyces paulus* NRRL 8115. Plos One 10(2015).

28. Losada, A. A. et al. Caboxamycin biosynthesis pathway and identification of novel benzoxazoles produced by cross-talk in *Streptomyces* sp NTK 937. Microb. Biotechnol. 10, 873–885 (2017).

29. Dosselaere, F. & Vanderleyden, J. A metabolic node in action: Chorismate-utilizing enzymes in microorganisms. Crit. Rev. Microbiol. 27, 75–131 (2001).

30. Maeda, H. & Dudareva, N. The shikimate pathway and aromatic amino acid biosynthesis in plants. Annu. Rev. Plant Biol. 63, 73–105 (2012).

31. Mavrodi, D. V., Blankenfeldt, W. & Thomashow, L. S. Phenazine compounds in fluorescent *Pseudomonas* spp. biosynthesis and regulation. Annu. Rev. Phytopathol. 44, 417–445 (2006).

32. Manthey, M. K., Pyne, S. G. & Truscott, R. J. Mechanism of reaction of 3-hydroxyanthranilic acid with molecular-oxygen. Biochim. Biophys. Acta. 1034, 207–212 (1990).

33. Parsons, J. F., Calabrese, K., Eisenstein, E. & Ladner, J. E. Structure and mechanism of *Pseudomonas aeruginosa* PhzD, an isochorismatase from the phenazine biosynthetic pathway. Biochemistry 42, 5684–5693 (2003).

34. Culbertson, J. E. & Toney, M. D. Expression and characterization of PhzE from *Pseudomonas aeruginosa* PAO1: aminodeoxyisochorismate synthase involved in pyocyanin and phenazine-1-carboxylate production. Biochim. Biophys. Acta. 1834, 240–246 (2013).

35. Muraki, T., Taki, M., Hasegawa, Y., Iwaki, H. & Lau, P. C. Prokaryotic homologs of the eukaryotic 3-hydroxyanthranilate 3,4-dioxygenase and 2-amino-3-carboxymuconate-6-semialdehyde decarboxylase in the 2-nitrobenzoate degradation pathway of *Pseudomonas fluorescens* strain KU-7. Appl. Environ. Microb. 69, 1564–1572 (2003).

36. Zhou, Y. et al. Determining the extremes of the cellular NAD(H) level by using an *Escherichia coli* NAD^+^-auxotrophic mutant. Appl. Environ. Microb. 77, 6133–6140 (2011).

37. Feng, J. et al. Bioretrosynthesis of functionalized N-heterocycles from glucose via one-pot tandem collaborations of designed microbes. Adv. Sci. 7, 2001188 (2020).

38. Zhang, L., Cao, Y., Tong, J. & Xu, Y. An alkylpyrazine synthesis mechanism involving l-threonine-3-dehydrogenase describes the production of 2,5-dimethylpyrazine and 2,3,5-trimethylpyrazine by *Bacillus subtilis*. Appl. Environ. Microb. 85 (2019).

39. Gao, H., Tiwari, M. K., Kang, Y. C. & Lee, J. K. Characterization of H_2_O-forming NADH oxidase from *Streptococcus pyogenes* and its application in L-rare sugar production. Bioorg. Med. Chem. Lett. 22, 1931–1935 (2012).

40. Mutti, F. G., Knaus, T., Scrutton, N. S., Breuer, M. & Turner, N.J. Conversion of alcohols to enantiopure amines through dual-enzyme hydrogen-borrowing cascades. Science 349, 1525–1529 (2015).

41. Houwman, J. A., Knaus, T., Costa, M. & Mutti, F. G. Efficient synthesis of enantiopure amines from alcohols using resting *Escherichia coli* cells and ammonia. Green Chem. 21, 3846–3857 (2019).

42. Thompson, M. P. & Turner, N.J. Two-enzyme hydrogen-borrowing amination of alcohols enabled by a cofactor-switched alcohol dehydrogenase. Chemcatchem 9, 3833–3836 (2017).

43. Pushpanath, A., Siirola, E., Bornadel, A., Woodlock, D. & Schell, U. Understanding and overcoming the limitations of *Bacillus badius* and *Caldalkalibacillus thermarum* amine dehydrogenases for biocatalytic reductive amination. ACS Catal. 7, 3204–3209 (2017).

44. Knaggs, A. R. The biosynthesis of shikimate metabolites. Nat. Prod. Rep. 18, 334–355 (2001).

## References

45. Sambrook, J. & Russell, D. W. Molecular Cloning: A Laboratory Manual, Third Edition; Cold Spring Harbor Laboratory Press: Cold Spring Harbor, New York (2001). Foundation, Norwich (2000).

46. Kieser, T., Bibb, M. J., Buttner, M. J., Chater, K. F. & Hopwood, D. A. Practical streptomyces genetics. John Innes Foundation, Norwich (2000).

47. Marshall, V. P., Little, M. S. & Johnson, L. E. A New Process and Organism for the Fermentation Production of Volonomycin. J. Antibiot. 34, 902–904 (1981).

48. Fernandez, E. et al. Identification of two genes from *Streptomyces argillaceus* encoding glycosyltransferases involved in transfer of a disaccharide during biosynthesis of the antitumor drug mithramycin. J. Bacteriol. 180, 4929–4937 (1998).

49. Li, P. et al. An efficient blue-white screening based gene inactivation system for *Streptomyces*. Appl. Microbiol. Biot. 99, 1923–1933 (2015).

50. Brumaghim, J. L., Li, Y., Henle, E. & Linn, S. Effects of hydrogen peroxide upon nicotinamide nucleotide metabolism in *Escherichia coli*: Changes in enzyme levels and nicotinamide nucleotide pools and studies of the oxidation of NAD(P)H by Fe(III). J. Biol. Chem. 278, 42495–42504 (2003).

51. Dawood, A. W., de Souza, R. O. & Bornscheuer, U. T. Asymmetric Synthesis of Chiral Halogenated Amines using Amine Transaminases. Chemcatchem 10, 951–955 (2018).

