## Supplemental Information for "Construction of a third NAD^+^ *de novo* biosynthesis pathway"

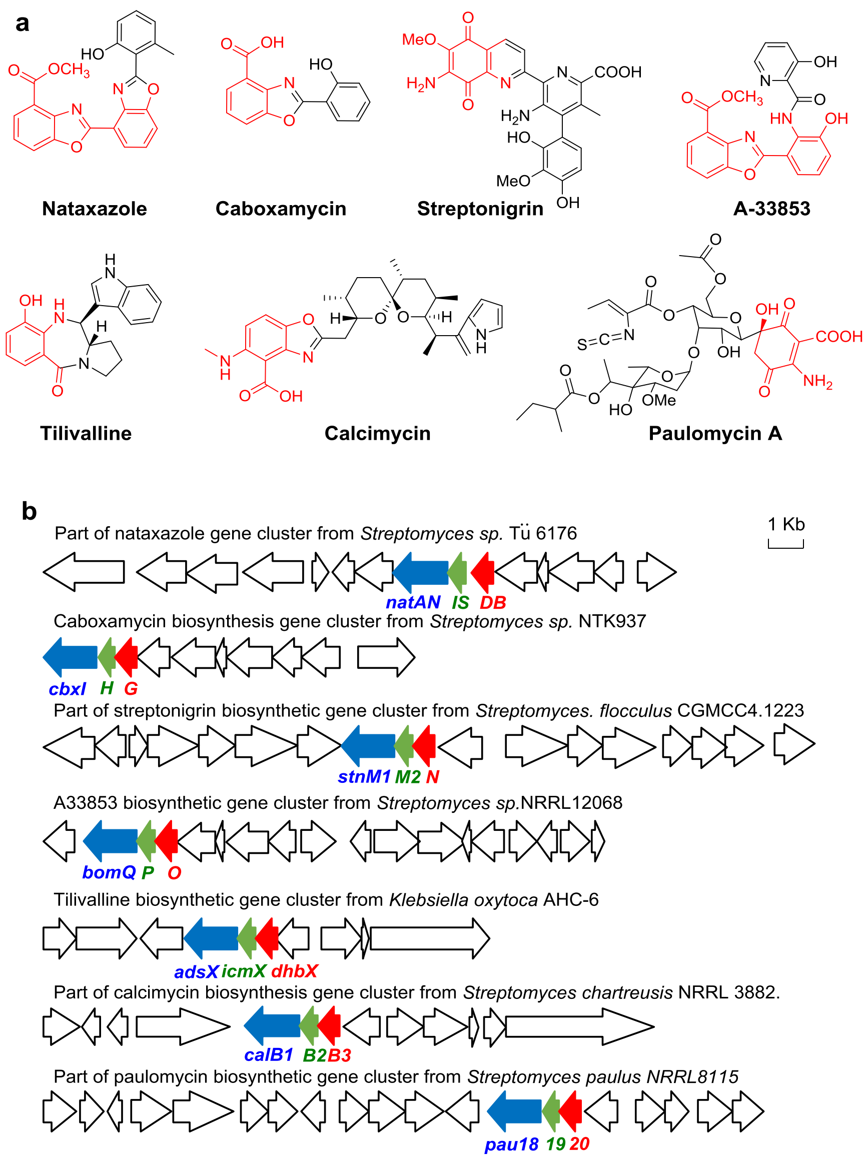


**Supplementary Fig. 1. Natural products with 3-HAA derived substructures and their biosynthetic gene clusters.** The three genes encoding ADIC synthase (blue), DHHA synthase (green), and DHHA dehydrogenase (red) are labeled in each cluster. DHHA dehydrogenases were proposed in these biosynthetic gene clusters but were not characterized biochemically until this study.

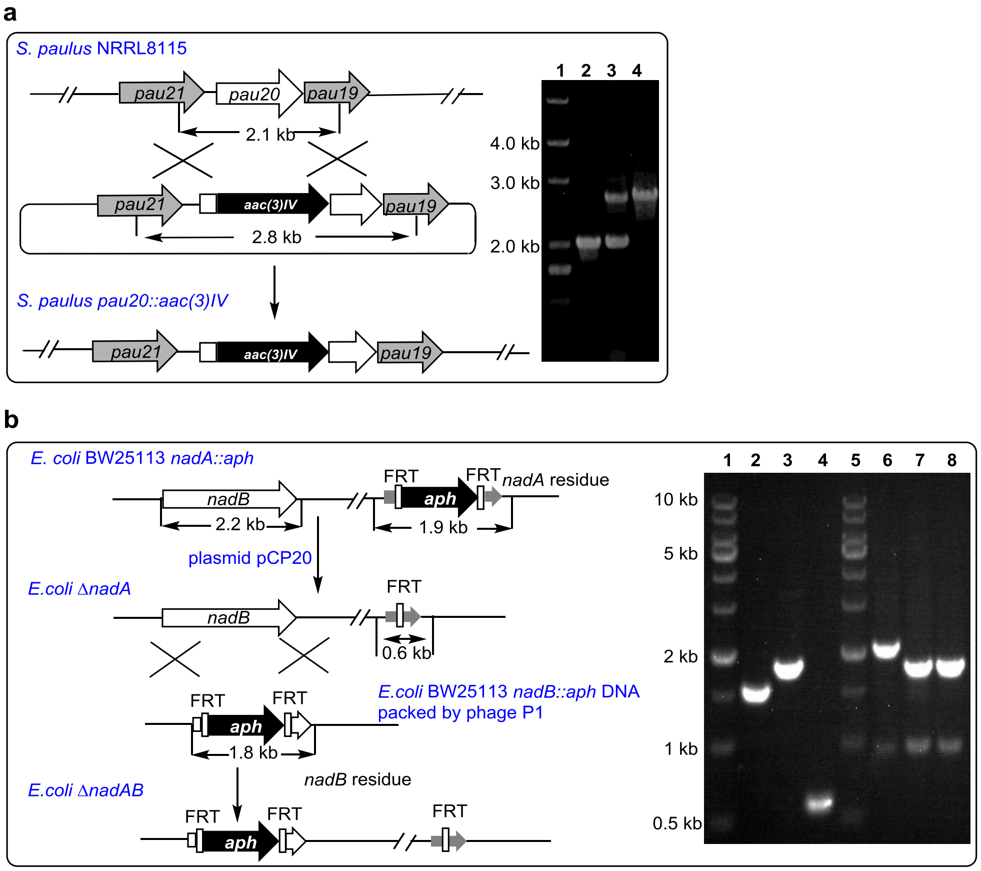


**Supplementary Fig. 2. Construction of *S. paulus pau20::aac(3)IV*, *E. coli* Δ*nadAB*, and *S. cerevisiae* Δ*BNA2*. a,** Illustration and PCR verification of *S. paulus pau20::aac(3)IV* construction: Lane 1, DNA marker; lane 2, *S. paulus* NRRL8115; lane 3, single crossover mutant; lane 4, *S. paulus pau20::aac(3)IV.* **b,** Illustration and PCR verification of *E. coli* Δ*nadAB* construction: Lane 1 and 5, DNA marker; lane 2 and 6, *E. coli* BW25113; lane 3 and 7, *E. coli* BW25113 *nadA::aph*, lane 4 and 8, *E. coli* Δ*nadAB.* For lane 2-4 and lane 6-8, the PCR reactions were performed with primers nadA-F/nadA-R and nadB-F/nadB-R, respectively.

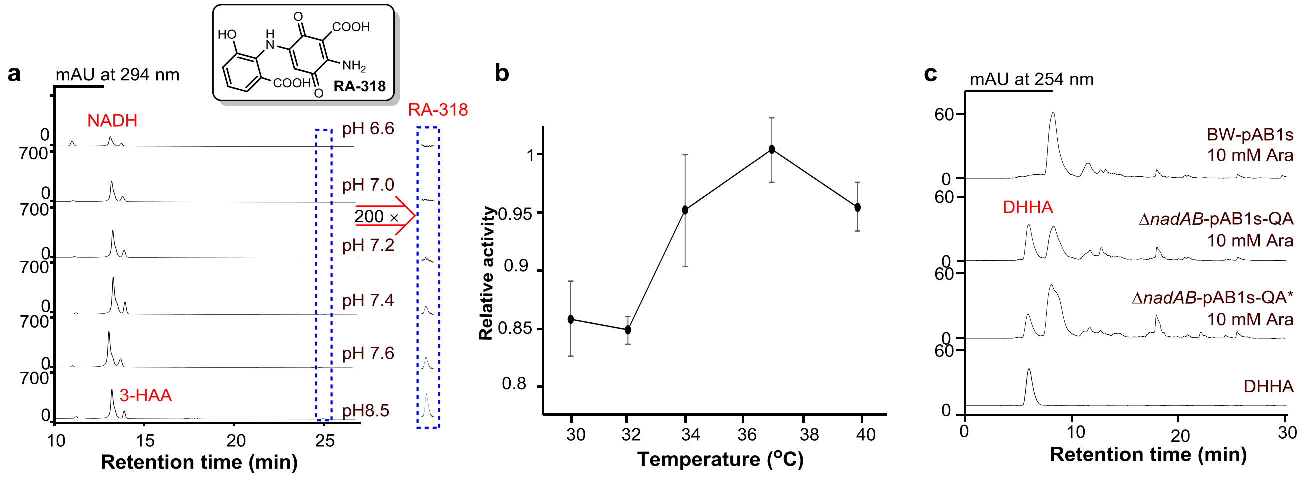


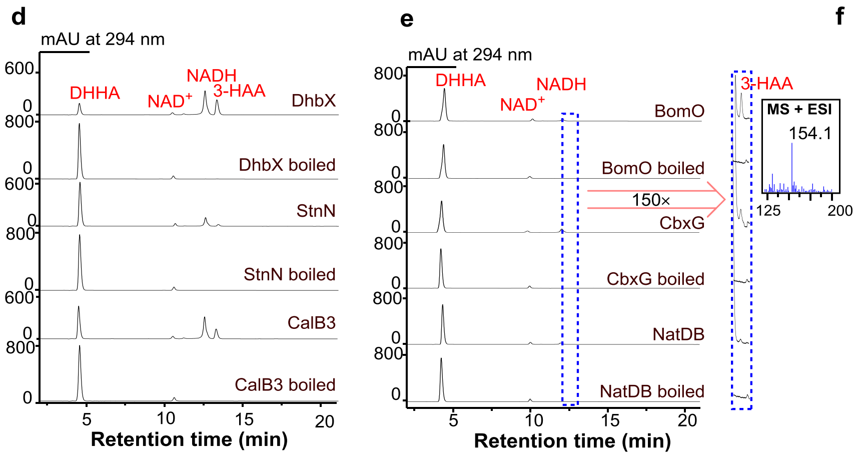


**f**

**Kinetic parameters of DHHA dehydrogenases**

| Toward NAD$^+$ | $K_m$ [mM] | $k_{cat}$ [min$^{-1}$] | $k_{cat}$/ $K_m$ [min$^{-1}$ mM$^{-1}$] |
|---|---|---|---|
| Pau20 | 0.40 ± 0.025 | 25.42 ± 0.72 | 63.55 ± 2.17 |
| CalB3 | 0.088 ± 0.017 | 11.06 ± 0.27 | 125.68 ± 21.21 |
| DhbX | 0.18 ± 0.054 | 50.44 ± 4.2 | 280.22 ± 60.73 |
| StnN | 0.33 ± 0.017 | 8.74 ± 0.14 | 26.48 ± 0.94 |
| **Toward DHHA** | $K_m$ [mM] | $k_{cat}$ [min$^{-1}$] | $k_{cat}$/ $K_m$ [min$^{-1}$ mM$^{-1}$] |
| Pau20 | 5.18 ± 0.55 | 33.00 ± 2.26 | 6.37 ± 0.09 |
| CalB3 | 2.96 ± 0.41 | 13.68 ± 1.10 | 4.62 ± 0.27 |
| DhbX | 1.69 ± 0.33 | 96.28 ± 11.08 | 56.97 ± 4.57 |
| StnN | 2.30 ± 0.31 | 12.84 ± 1.18 | 5.58 ± 0.24 |

**Supplementary Fig. 3. Enzymatic studies to identify more efficient DHHA dehydrogenases. a,** Pau20 reactions in buffers with varied pH values: Pau20 assays were carried out at 37 °C for 30 min in buffers with pH values ranging from 6.6 to 8.5 unless specified. As previously reported, compound RA-318 was the main spontaneously oxidized product of 3-HAA under the assay conditions; its structure was assigned by MS and NMR analyses (data not shown). **b,** Optimization of Pau20 reaction temperature: Pau20 assays were carried out in 200 mM phosphate buffer (pH 7.0) for 30 min with temperatures ranging from 30 to 40 °C. **c,** HPLC metabolite profiles of *E. coli* BW-pAB1s, Δ*nadAB*-pAB1s-QA and Δ*nadAB*-pAB1s-QA* fermentation broth to check the accumulations of DHHA in those strains during early stationary phase. **d,** Representative assays of DHHA dehydrogenases DhbX, StnN, and CalB3. Those assays were carried out at 37 °C for 2 hours in 200 mM phosphate buffer (pH 7.0). **e,** Representative assays of DHHA dehydrogenases BomO, CbxG, and NatDB. The production of 3-HAA was confirmed by LC-MS. Those assays were carried out at 37 °C for 2 hours in 200 mM phosphate buffer (pH 7.0). **f,** Steady-state kinetic parameters of DHHA dehydrogenases Pau20, DhbX, StnN, and CalB3 at 37 °C pH 7.0.

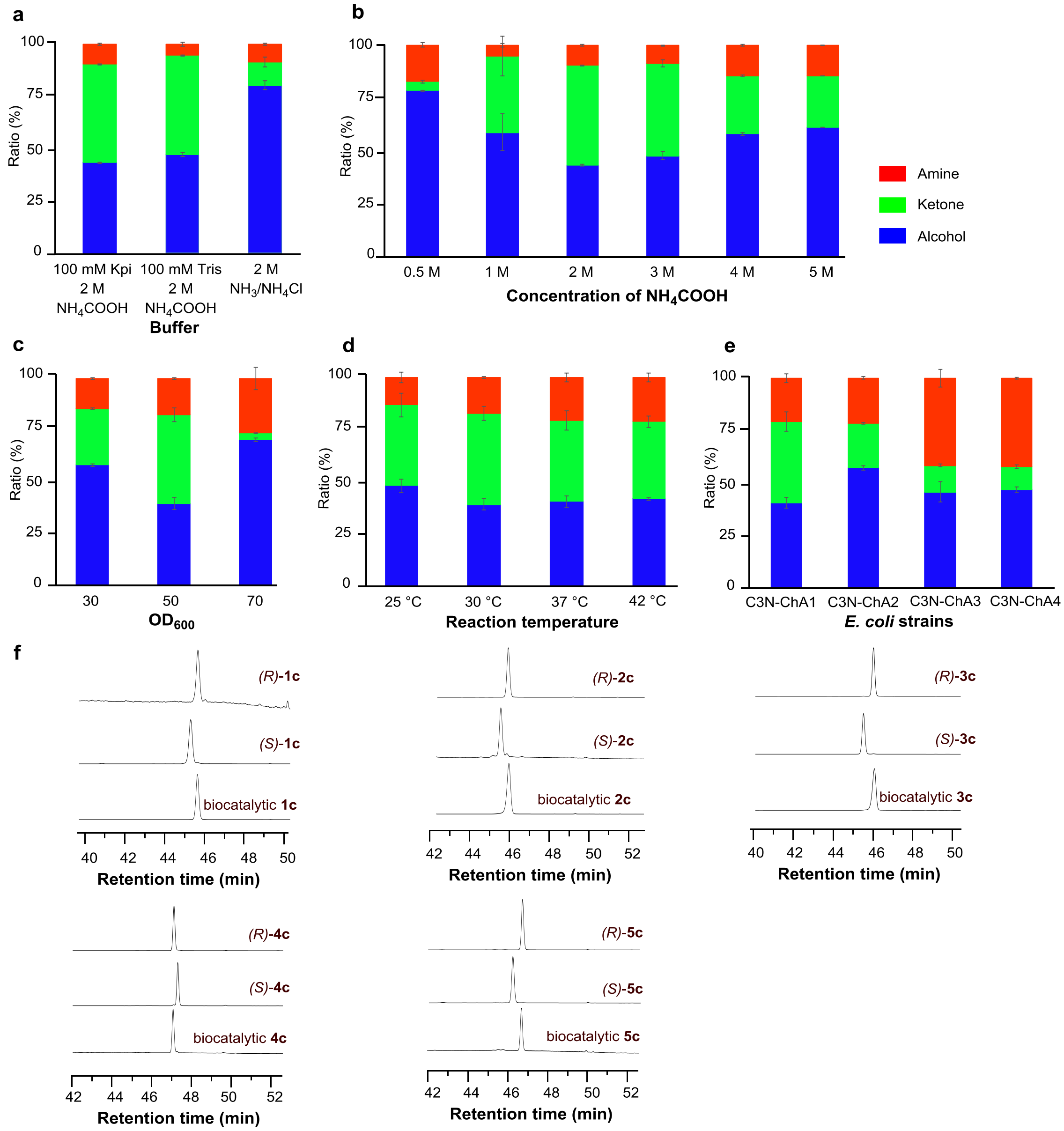


**Supplementary Fig. 4. Optimization of the bioamination procedure using *rac*-1a as a substrate and enantiomeric purity analysis of the chiral amines produced by the C3N pathway-based whole-cell systems. a,** Optimization of catalytic buffers (pH 8.5): 5 mM *rac*-**1a**, $OD_{600}$ = 30, under 30 °C, 230 rpm for 10 h in 1 mL reaction volume. All tested buffers contained 10% DMSO. **b,** Optimization of the concentration of $NH_4COOH$: 100 mM KPi buffer (pH 8.5) with 10% DMSO, 5 mM *rac*-**1a**, $OD_{600}$ = 30, under 30 °C, 230 rpm for 10 h in 1 mL reaction volume. **c,** Optimization of biomass ($OD_{600}$): 100 mM KPi buffer (pH 8.5) with 10% DMSO, 5 mM *rac*-**1a**, 4 M $NH_4COOH$, under 30 °C, 230 rpm for 10 h in 1 mL reaction volume. **d,** Optimization of reaction temperature: 100 mM KPi buffer (pH 8.5) with 10% DMSO, 5 mM *rac*-**1a**, 4 M $NH_4COOH$, $OD_{600}$ = 50, 230 rpm for 10 h in 1 mL reaction volume. **e,** Detection of the bioamination capacities of different engineering strains: 100 mM KPi buffer (pH 8.5) with 10% DMSO, 5 mM *rac*-**1a**, 4 M $NH_4COOH$, $OD_{600}$ = 50, under 37 °C, 230 rpm for 10 h in 1 mL reaction volume. **f,** HPLC analysis of the enantiomeric purities of the chiral amines derivatized by FDAA (Marfey's Reagent).

**Supplementary Table 1.** Bacterial strains and plasmids.

| **Strains or plasmids** | **Characteristics**[a] | **Reference or source** |
|---|---|---|
| ***Escherichia coli*** | | |
| JM 109 | General cloning host | Lab stock |
| BL21(DE3) | Host for protein expression | Novagen |
| ET12567/pUZ8002 | Strain for intergeneric conjugation | Invitrogen |
| BW25113 | Wild-type strain | Lab stock |
| BW25113 *nadA::aph* | BW25113 *nadA* mutant | 1 |
| BW25113 *nadB::aph* | BW25113 *nadB* mutant | 1 |
| BW25113 Δ*nadAB* | BW25113 *nadA & nadB* combined mutant | This work |
| BW-pXB1s-HAA | BW25113 harboring pXB1s-HAA | This work |
| BW-pXB1s | BW25113 harboring pXB1s | This work |
| BW-pAB1s | BW25113 harboring pAB1s | This work |
| Δ*nadAB*-pXB1s-QA | BW25113 Δ*nadAB* harboring pXB1s-QA | This work |
| Δ*nadAB*-pAB1s-QA | BW25113 Δ*nadAB* harboring pAB1s-QA | This work |
| Δ*nadAB*-pXB1s-QA* | BW25113 Δ*nadAB* harboring pXB1s-QA* | This work |
| Δ*nadAB*-pAB1s-QA* | BW25113 Δ*nadAB* harboring pAB1s-QA* | This work |
| BW25113(DE3) | Chassis cell for cell factory | Lab stock |
| DMP-Con | BW25113(DE3) harboring pAB1s and pRSF-*EcTdh-SpaNox* | This work |
| C3N-DMP | BW25113(DE3) harboring pAB1s-QA* and pRSF-*EcTdh-SpaNox* | This work |
| BW-ChA1-Con | BW25113(DE3) harboring pAB1s and pCDF-*TesADH-CalAmDH* | This work |
| BW-C3N-ChA1 | BW25113(DE3) harboring pAB1s-QA* and pCDF-*TesADH-CalAmDH* | This work |
| BW-C3N-ChA2 | BW-C3N-ChA1 harboring pACYC-*CalAmDH* | This work |
| BW-C3N-ChA3 | BW25113(DE3) harboring pAB1s-QA* and pRSF-*TesADH-CalAmDH* | This work |
| BW-C3N-ChA4 | BW-C3N-ChA3 harboring pACYC-*CalAmDH* | This work |
| BW-ChA3-Con | BW25113(DE3) harboring pAB1s and pRSF-*TesADH-CalAmDH* | This work |
| ***Streptomyces*** | | |
| *S. paulus* NRRL 8115 | Wild-type strain | NRRL |
| *S. paulus* NRRL 8115 *pau20::aac(3)IV* | *S. paulus* NRRL 8115 *pau20* mutant | This work |
| **Plasmids** | | |
| pET28a | protein production vector, $Kan^r$ | Novagen |
| pET28a-*pau20* | pET28a harboring *pau20*, $Kan^r$ | This work |
| pET28a-*dhbX* | pET28a harboring *dhbX*, $Kan^r$ | This work |
| pET28a-*calB3* | pET28a harboring *calB3*, $Kan^r$ | This work |

| | | |
|---|---|---|
| pET28a-*cbxG* | pET28a harboring *cbxG*, Kan$^r$ | This work |
| pET28a-*stnN* | pET28a harboring *stnN*, Kan$^r$ | This work |
| pET28a-*bomO* | pET28a harboring *bomO*, Kan$^r$ | This work |
| pET28a-*natDB* | pET28a harboring *natDB*, Kan$^r$ | This work |
| pAB1s (high-copy-number) | ColE1 origin, *ara*BAD promoter, Sm$^r$ | 2 |
| pXB1s (medium-copy-number) | p15A origin, *ara*BAD promoter, Sm$^r$ | 2 |
| pAB1s-HAA | pAB1s harboring *pau20*, *phzD* and *phzE* | This work |
| pXB1s-HAA | pXB1s harboring *pau20*, *phzD* and *phzE* | This work |
| pAB1s-QA | pAB1s harboring *nbaC*, *pau20*, *phzD* and *phzE* | This work |
| pXB1s-QA | pXB1s harboring *nbaC*, *pau20*, *phzD* and *phzE* | This work |
| pAB1s-QA* | pAB1s harboring *nbaC*, *dhbX*, *phzD* and *phzE* | This work |
| pXB1s-QA* | pXB1s harboring *nbaC*, *dhbX*, *phzD* and *phzE* | This work |
| pACYCduet-1 | p15 origin, T7 promoter, Cm$^r$ | Novagen |
| pCDFduet-1a | CDF origin, T7 promoter, Amp$^r$ | Novagen |
| pRSFduet-1 | RSF origin, T7 promoter, Kan$^r$ | Novagen |
| pRSF-*EcTdh-SpaNox* | pRSFduet-1 harboring *EcTdh* and *SpaNox* | This work |
| pET28a-*TesADH* | pET28a harboring *TesADH*, Kan$^r$ | This work |
| pET28a-*CalAmDH* | pET28a harboring *CalAmDH*, Kan$^r$ | This work |
| pACYC-*CalAmDH* | pACYCduet-1 carrying *CaLAmDH*, Cm$^r$ | This work |
| pCDF-*TesADH* | pCDFduet-1a harboring *TesADH*, Amp$^r$ | This work |
| pCDF-*TesADH-CalAmDH* | pCDFduet-1a harboring *TesADH* and *CaLAmDH*, Amp$^r$ | This work |
| pRSF-*TesADH-CalAmDH* | pRSFduet-1a harboring *TesADH* and *CaLAmDH*, Amp$^r$ | This work |

[a]: Kan$^r$, kanamycin resistance; Sm$^r$, streptomycin resistance; Amp$^r$, ampicillin resistance; Cm$^r$, chlorampenicol resistance.

**Supplementary Table 2. Oligonucleotides used in this work.**

| Name | Sequence(5′→3′) |
|---|---|
| pau20-s2 | cagtcgattggctgacaattgattccgctcggcaggttcg |
| pau18R | cttgctagcagatgtcaattgatccgggccatcatcttcagt |
| pau22-R | taaaacgacggccagtgaattccatatggcgtcgcgccaccggcc |
| pau20-R | atcccttaacgtgagcctaggcagtcgtcctcggtgagttccag |
| pauN10ES | aaaactgcagcatatgggcacagccaattccgac |
| pauN10ER | cgggatccctggatgggcgtgagcgtc |
| pHAApau20-F | caggaggaattaaccatgggcacagccaattccg |
| pHAApau20-R | ctcctctttctctagacatatgctagcggcccagggtcgcgc |
| pHAAphzDE-F | catatgtctagagaaagagg |
| pHAAphzDE-R | accgagctcaccgaattcggatccttatgggcgacg |
| NbaC-F | gctaacaggaggaattaaccatgatgtttacctttggtaaac |
| NbaC-R | tcggaattggctgtgcccatctagtatttctcctctttctctagaggatccttacggctgatcac |
| nadA-F | tcaggcatcctcaatttc |
| nadA-R | ggcatacagctgaatctg |
| nadB-F | aacatcgcattatctgtg |
| nadB-R | gcgtagtgctgccagagc |
| T7 | taatacgactcactatagg |
| Cal-P23R | gatgtaggtgttccacaggcaaaaaacccctcaagacccg |
| CALRS-F | aagaaggagatatacatatgtctaccgtgacctttg |
| CALRS-R | taccagactcgagggtacttagcgacgaacgcgccat |
| Cal-his-R | agtgcggccgcaagcttgtcgacttagcgacgaacgcgccatt |

### References


1. Baba, T. *et al.* Construction of *Escherichia coli* K-12 in-frame, single-gene knockout mutants: the Keio collection. *Mol. Syst. Biol.* **2** (2006).
2. Cui, Q., Zhou, F., Liu, W. & Tao, Y. Avermectin biosynthesis: stable functional expression of branched chain alpha-keto acid dehydrogenase complex from *Streptomyces avermitilis* in *Escherichia coli* by selectively regulating individual subunit gene expression. *Biotechnol. Lett.* **39**, 1567-1574 (2017).
